# Thermodynamically-consistent, reduced models of gene regulatory networks

**DOI:** 10.1101/2023.11.13.566770

**Authors:** Michael Pan, Peter J. Gawthrop, Matthew Faria, Stuart T. Johnston

## Abstract

Synthetic biology aims to engineer novel functionalities into biological systems. While the approach has to date been predominantly applied to single cells, a richer set of biological phenomena can be engineered by applying synthetic biology to cell populations. To rationally design cell populations, we require mathematical models that link between intracellular biochemistry and intercellular interactions. In this study, we develop a kinetic model of gene expression that is suitable for incorporation into agent-based models of cell populations. To be scalable to large cell populations, models of gene expression should be both computationally efficient and compliant with the laws of physics. We satisfy the first requirement by applying a model reduction scheme to translation, and the second requirement by formulating models using bond graphs. Our reduced model is significantly faster to simulate than the full model, and faithfully reproduces important behaviours of the full model. We couple separate models of gene expression to build models of the toggle switch and repressilator. With these models, we explore the effects of resource availability and cell-to-cell heterogeneity on circuit behaviour. The modelling approaches developed in this study are a bridge towards engineering collective cell behaviours such as synchronisation and division of labour.

## 1 Introduction

Synthetic biology is the engineering of novel functionalities into biological systems. This enables biological solutions to problems that are difficult or impossible to solve with conventional engineering technologies, such as the detection and breakdown of toxic environmental pollutants and the production of biological therapeutics [1, 2]. Synthetic biology has been applied with great success to single cells, making use of synthetic genes as the fundamental building block. The field first gained attention through early biological implementations of two simple genetic circuits; the genetic *toggle switch*, which is capable of flipping between two stable states [3]; and the *repressilator*, a negative feedback loop composed of three repressor genes capable of oscillations [4]. Since then, synthetic biology has been used to design more complex systems such as biosensors and cell computers [5, 6]. To scale up from simple to complex systems, synthetic biology employs a modular approach from engineering. Thus, standard modules (genetic constructs that have been designed according to specifications) can be re-used as part of a more complex system. With this approach, even relatively simple genetic circuits can be assembled into biological logic gates, and ultimately biological devices capable of making complex decisions [6].

There are several known limitations to single-cell synthetic biology. Firstly, there may be undesired cross-talk between synthetic genes, leading to unpredictable function [7]. Secondly, the newly inserted genes consume the same resources (energy, ribosomes and amino acids) as other pathways in the cell [8]. Thus, a resource-intensive gene may compete with the cell’s endogenous pathways, leading to poor growth [9]. Finally, it cannot be assumed that upstream modules are completely isolated from the effects of downstream modules. In general, downstream pathways will influence and in some cases disrupt the behaviour of upstream pathways. This is known as retroactivity, and can lead to poor modularity even in the absence of cross-talk [10].

To address the known limitations of single-cell synthetic biology, synthetic biology of cell populations and consortia has been proposed [9, 11]. Synthetic modules can be inserted into different cells of a consortium, physically isolating them from each other [5]. This reduces the risk of cross-talk, competition and retroactivity between modules. Furthermore, it is possible to design population-level behaviours that would not be possible within homogeneous populations. For example, the quorum sensing system in bacteria is a frequent target of synthetic biology [12]. This system senses local cell density via the accumulation of signalling molecules, and can be engineered to reduce noise and synchronise oscillations [9]. It is also possible to engineer cell populations to form various spatial patterns when grown on media [13].

In spite of the clear potential of cell populations for synthetic biology, it is still an emerging field compared to single-cell synthetic biology. A major challenge is predicting how changes at the single-cell level affect behaviour at the population scale. Because it is infeasible to physically test every possible implementation of a synthetic biological system, synthetic biologists use design-build-test-learn cycles to optimise performance with limited time and resources [1]. In this context, mathematical models are essential to ruling out infeasible designs prior to implementation. Mathematical models have been widely used to describe cellular biochemistry (systems biology) and interactions between cells (agent-based models) [14]. While it is possible to model cell populations without agent-based models, coupling systems biology models to agent-based models is essential to understanding phenomena such as cell heterogeneity, synchronisation and cell migration. Such population-level dynamics pose a unique set of challenges for modelling engineered cell populations. In particular, cell populations can easily consist of millions of cells [15]. Thus, while complex models of gene regulatory networks are sufficient for single-cell synthetic biology, computational efficiency becomes a major consideration in engineering cell populations [16].

Because synthetic biology is focussed on engineering gene circuits, modelling gene regulatory networks is a key point of interest. The landmark studies implementing the toggle switch and repressilator used simple ordinary differential equation models of gene regulation, ignoring the dynamics of transcription and translation [3, 4]. This approach has been generalised and expanded upon to model large-scale gene regulatory networks [17]. However, the above models do not explicitly model transcription and translation, nor do they account for ribosome and energy availability. More recent models have sought to incorporate this level of detail. Whole-cell models of *Mycoplasma genitalium* and *Escherichia coli* account for the use of ribosomes, amino acids and energy in translation [18, 19]. Weiße et al. [20] take a simpler approach, using generic molecules for energy and monomers as variables in a differential equation model of transcription and translation. There are also more detailed models of translation that account for the position of ribosomes on mRNA and can predict the occurrence of ribosomal ‘traffic jams’. These include the stochastic totally asymmetric exclusion process (TASEP) and its deterministic approximation the ribosome flow model (RFM) [21]. However, a consideration for the thermodynamics of gene expression has for the most part been lacking. Many models completely ignore the energy molecule adenosine triphosphate (ATP) and therefore do not account for the dependence of expression dynamics on energy. This is surprising given that translation consumes four ATP molecules per amino acid [22], making it one of the most energy-consuming processes in a cell (30-75%) [18, 23]. If we are to adequately model gene regulatory networks for the synthetic biology of cell populations, a new modelling approach is needed that accounts for energy while also resulting in models simple enough to be incorporated into large cell populations.

In this study, we propose a new, simplified, and thermodynamically consistent model of gene expression. The model satisfies three key criteria: it is (i) computationally feasible to simulate; (ii) dependent on the availability of energy and ribosomes; and (iii) thermodynamically consistent. We primarily focus on translation as it is a major consumer of energy. A model reduction scheme is developed to simplify the chains of reactions involved in translation elongation. This ensures that the model is simple but biochemically plausible, i.e. it is a limiting case of a more complex reaction network. To ensure thermodynamic consistency, we use the bond graph approach, which is a powerful, modular framework for modelling biological systems in a physically consistent manner [24]. Previous work has shown that thermodynamic constraints automatically fall out from the parameterisation inherent to bond graph models, and that the approach is scalable to large biochemical networks [25–29]. Furthermore, bond graphs provide a natural methodology for accounting for retroactivity [30]. We exploit the graphical, modular nature of bond graphs to couple models of gene expression together, enabling the construction of common synthetic biology motifs such as the toggle switch and repressilator.

In Section 2, we introduce the bond graph formalism, which is then used to construct thermodynamically consistent models of gene regulatory networks. A model reduction scheme for translation is developed to ensure the model is simple but still retains important features of a more detailed model (§ 2.4). By coupling models of gene expression together, models of synthetic circuits are developed (§ 2.6). We first compare the full and reduced models, and show that their steady-state behaviours match well (§ 3.1). We next show that the bond graph model is able to reproduce experimentally observed behaviours in common synthetic biology motifs. In § 3.2–3.3, we investigate the effects of energy and ribosome availability on synthetic circuits. We then examine the potential effects of parameter heterogeneity in synthetic circuits (§ 3.4). Finally, Section 4 contains a discussion of the results and concluding remarks.

## 2 Methods

### 2.1 Overview

The gene is a key building block of synthetic biology. Here, we consider two key processes in gene expression: *transcription*, which produces mRNA from a DNA template; and *translation*, which produces protein from an mRNA template (Figure 1). This section will detail the methods required to model gene expression in a thermodynamic framework. Inspired by Weiße et al. [20], we adopt a simplistic approach for modelling these processes. In particular, we use generic species to model cell resources rather than accounting for all their constituent biomolecules. Thus, both energy (*A*) and ribosomes (*R*) are represented as individual species. Figure 1 depicts an overview of the model. It is assumed that the reactions take place in a constant volume, i.e. in the absence of cell expansion.

**Figure 1:**
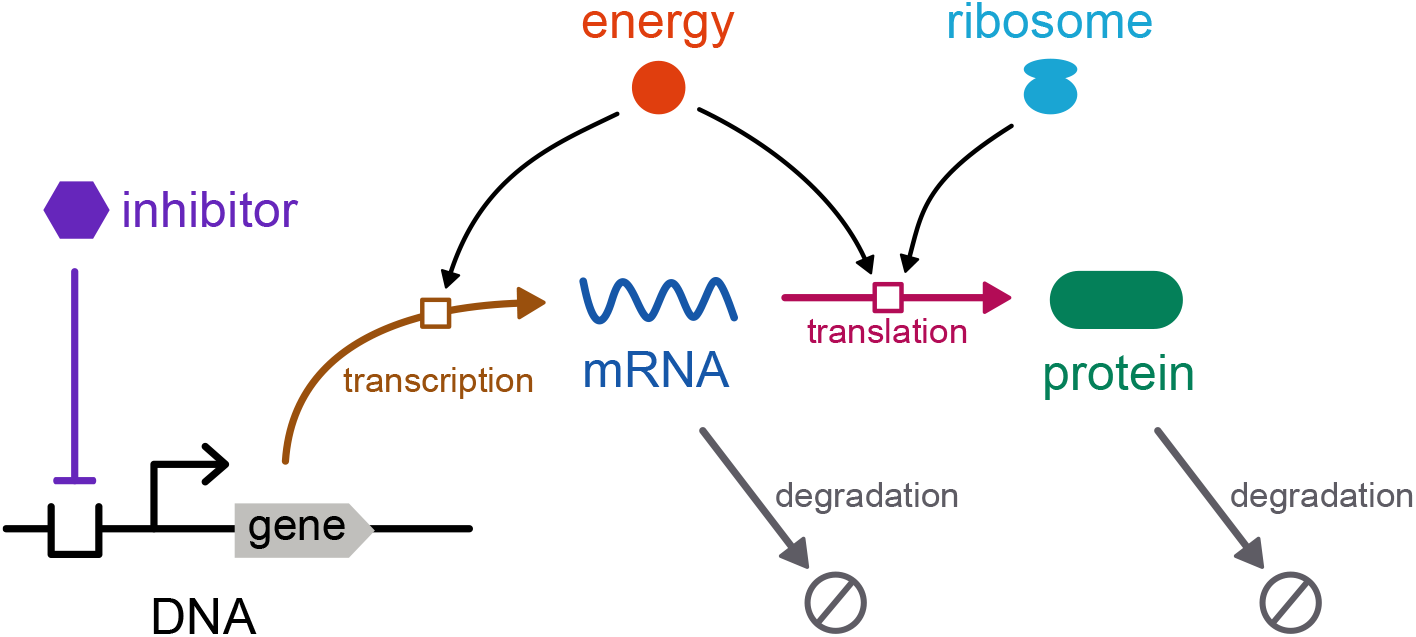
Schematic of gene expression model.

We first introduce the bond graph approach for modelling biochemical systems, and explain how it ensures adherence to the laws of thermodynamics as well as its relation to kinetic modelling (§ 2.2). We then develop bond graph models of transcription (§ 2.3) and translation (§ 2.4). To ensure that the model of translation is computationally tractable, we apply a model reduction scheme (§ 2.4.2). We combine the models of transcription and translation into a model of gene expression (§ 2.5) and detail how models of gene expression can be composed into models of common synthetic biology motifs (§ 2.6).

### 2.2 Bond graph modelling

The bond graph is a powerful and general methodology for modelling biological systems while adhering to the laws of physics and thermodynamics. We outline the application of bond graphs to biochemistry specifically here; the reader is referred to Gawthrop and Pan [24] for more general applications to biology.

As an illustrative example, the bond graph of the reaction network

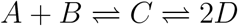

is given in Figure 2, with details explained in the following paragraphs.

**Figure 2:**
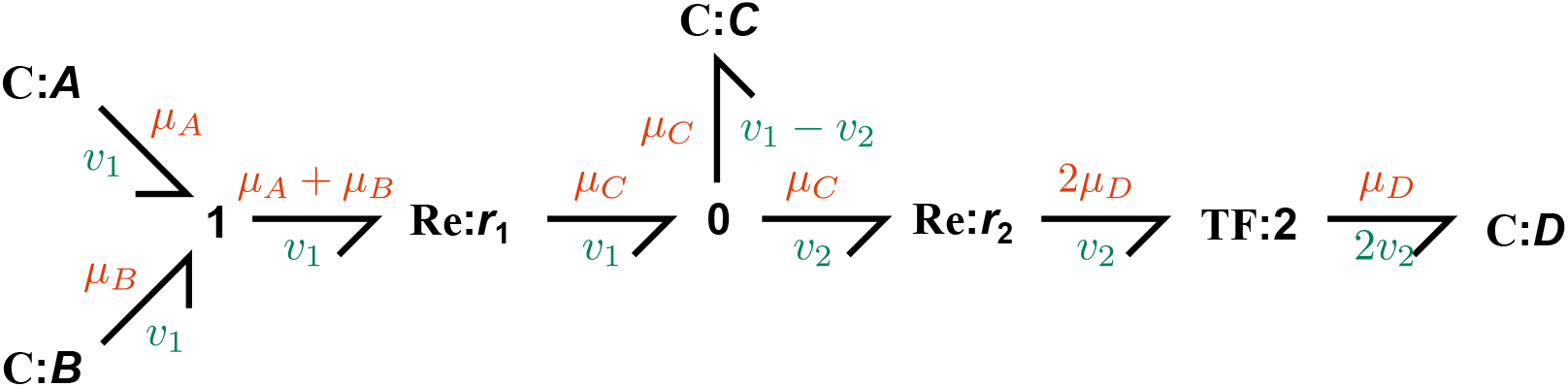
The bond graph of the reaction network. *A* + *B* ⇌ *C* ⇌ 2*D*. Each bond is labelled with a potential *μ* and a flow *v*. The components follow the notation *component type*: *name*.

Bond graphs are a generalisation of directed graphs, with physical *components* as the nodes and *bonds* as the edges. Bonds are the fundamental unit of connection in bond graphs, and are depicted using harpoons. Each bond carries two physical covariables: the chemical potential *μ* [J/molecule] and the molecular flow rate *v* [molecules/s]. Since the product of *μ* and *v* is power [J/s], bond graphs track the movement of energy between components and ensure thermodynamic consistency.

#### 2.2.1 Chemical species

Chemical species are labelled as **C** components because they store energy in their concentration. This can be thought of in the context of diffusion: particles in a region of higher concentration have higher potentials and therefore tend to move to regions of lower concentration. In dilute solutions, the chemical potential *μ*_*i*_ of a chemical species *i* is given by

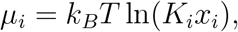

where *k*_*B*_ = 1.381*×*10^*−*23^ J*/*K is the Boltzmann constant, *T* [K] is the absolute temperature, *K*_*i*_ [molecules^*−*1^] is a thermodynamic parameter and 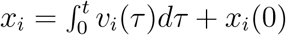 is the molar amount of species, defined as the integral of molecular flow [31]. For convenience, we define the dimensionless chemical potential as

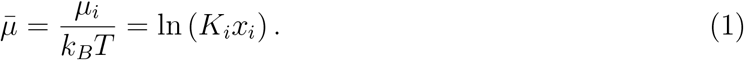

In some cases, species are fixed at constant concentrations due to an external supply. These are represented as **Se** components whose concentrations remain constant.

#### 2.2.2 Reactions

Reactions convert chemical species into other chemical species, and are labelled as **Re** components. Due to the principle of microscopic reversibility, all reactions are reversible in reality and therefore are modelled as reversible reactions [32]. Assuming mass-action kinetics, the flux through a reaction *j* is given by the Marcelin-de Donder equation

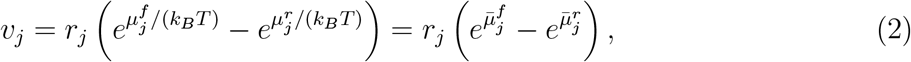

where *r*_*j*_ is the reaction rate parameter, 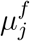 is the sum of reactant chemical potentials and 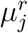 is the sum of product chemical potentials [25]. The bar notation indicates dimensionless chemical potentials.

#### 2.2.3 Network topology

When a species is involved in multiple reactions, they are connected to **0** junctions. This ensures that the species accumulate at a rate equal to the sum of contributions from each reaction (mass balance). The junction also ensures that the species contributes the same chemical potential to each reaction. For example, the **0** junction in Figure 2 ensures that the flow of the bond into the species *C* is equal to the difference between flows of the other two connected bonds 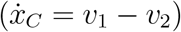.

When either side of a reaction involves multiple species, the species are connected to a **1** junction prior to the reaction. This junction ensures that all species are consumed or produced at the same rate, and that the contributions of the chemical potentials are summed appropriately. For example, the **1** junction in Figure 2 ensures that the chemical potential on the outgoing bond, *μ*_*A*_ + *μ*_*B*_, is equal to the sum of chemical potentials of incoming bonds.

#### 2.2.4 Stoichiometry

If a species contributes multiple stoichiometry to a reaction, it is connected through a **TF** component. This component encodes the relationship

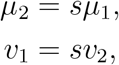

where *s* is the stoichiometry, the subscript 1 indicates the bond closer to the species and the subscript 2 indicates the bond closer to the reaction. The **TF** component ensures that the chemical potential is counted multiple times prior to calculating the contribution of a species to a reaction, and that flux is scaled up by the stoichiometric constant.

#### 2.2.5 Relationship to kinetic modelling

A general methodology for constructing bond graphs of biochemical systems is given in Gawthrop and Crampin [25]. Using this approach, the potential balance constraints of the **1** junctions give rise to the equations

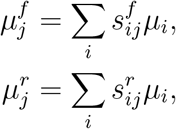

where 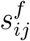 is the stoichiometric contribution of species *i* to the forward side of reaction *j*, and similarly for 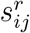 on the reverse side.

By using Eqs. (1)–(2), we find that

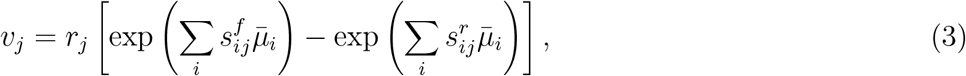

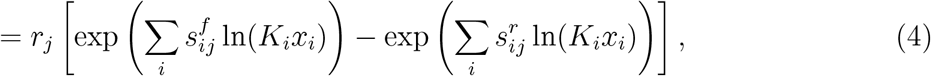

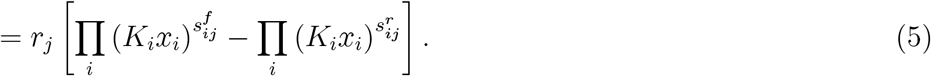

This is the well-known law of mass action

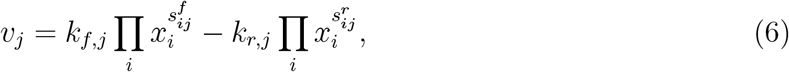

where the forward and reverse kinetic parameters are defined as

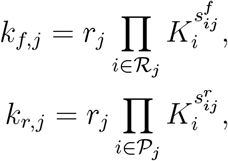

where ℛ_*j*_ is the set of all reactants and 𝒫_*j*_ the set of all products in reaction *j*. Finally, the **0** junctions ensure mass balance, thus

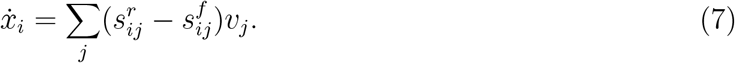

Eqs. (5) and (7) give rise to a set of differential equations for the biochemical system. Thus, the bond graph in Figure 2 encodes the differential equations

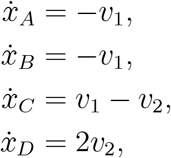

Where

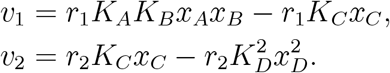

### 2.3 Transcription

Following Weiße et al. [20], we model transcription as an enzyme-catalysed reaction with the energy (*A*) as the substrate and mRNA (*M*) as the product, under the effect of an inhibitor *I* (such as a transcription factor). We assume non-competitive inhibition, so the net rate of reaction under the quasi-steady-state approximation is

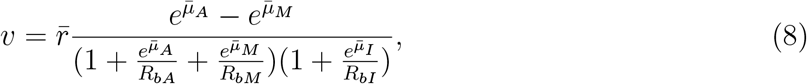

Where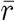 is a rate parameter and *R*_*bA*_, *R*_*bM*_ and *R*_*bI*_ are affinity parameters (see § A.1 of the electronic supplementary material for further detail). Thus, Eq. (8) is represented as the three-port **Tc** component presented in Figure 3, which is a bond graph representation of transcription that reproduces the energy-saturating behaviour in Weiße et al. [20]. While we only consider the inhibition of transcription in this study, promoters can be modelled by employing a similar approach on an enzyme activation reaction scheme [33].

**Figure 3:**
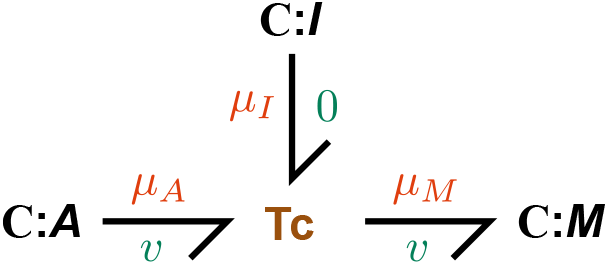
Bond graph model of transcription. The **Tc** component has the rate law described in Eq. (8).

### 2.4 Translation

#### 2.4.1 Full model

Because we explicitly model the use of energy and ribosomes in translation, a greater level of detail is required compared to the modelling of transcription. We first describe a detailed model, which is used as the basis for deriving a reduced model that is necessary for modelling large cell populations. Our model of translation considers the following processes:

1. **Binding** of ribosomes (*R*) to mRNA (*M*) to create a complex (*C*_0_).
2. **Elongation**, the movement of ribosomes along the mRNA. For every site that the ribosome moves, a molecule of energy (*A*) is consumed, increasing the length of the nascent protein.
3. **Termination**, the release of ribosome and the completed protein (*P*) from the mRNA.

The reaction scheme that corresponds to these processes is:

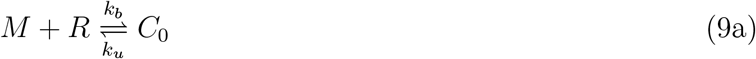

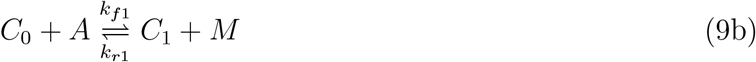

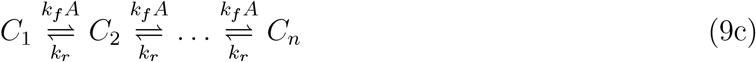

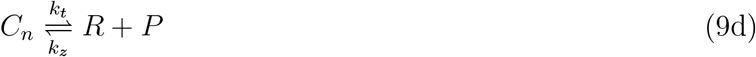

where *n* is the number of ATP molecules required to synthesise the protein. Consistent with the TASEP model [21], we assume that multiple ribosomes can bind to the same transcript, thus the mRNA molecule is released after the first elongation step (Eq. (9b)). However, for simplicity, we assume that apart from the first binding site, ribosomes do not obstruct each other, so that the complexes may be treated independently. While both binding and termination kinetics are dependent on the mRNA sequence [34, 35], we assume for simplicity that both initiation and termination dynamics are independent of sequence. A bond graph of the full model is shown in Figure 4a.

**Figure 4:**
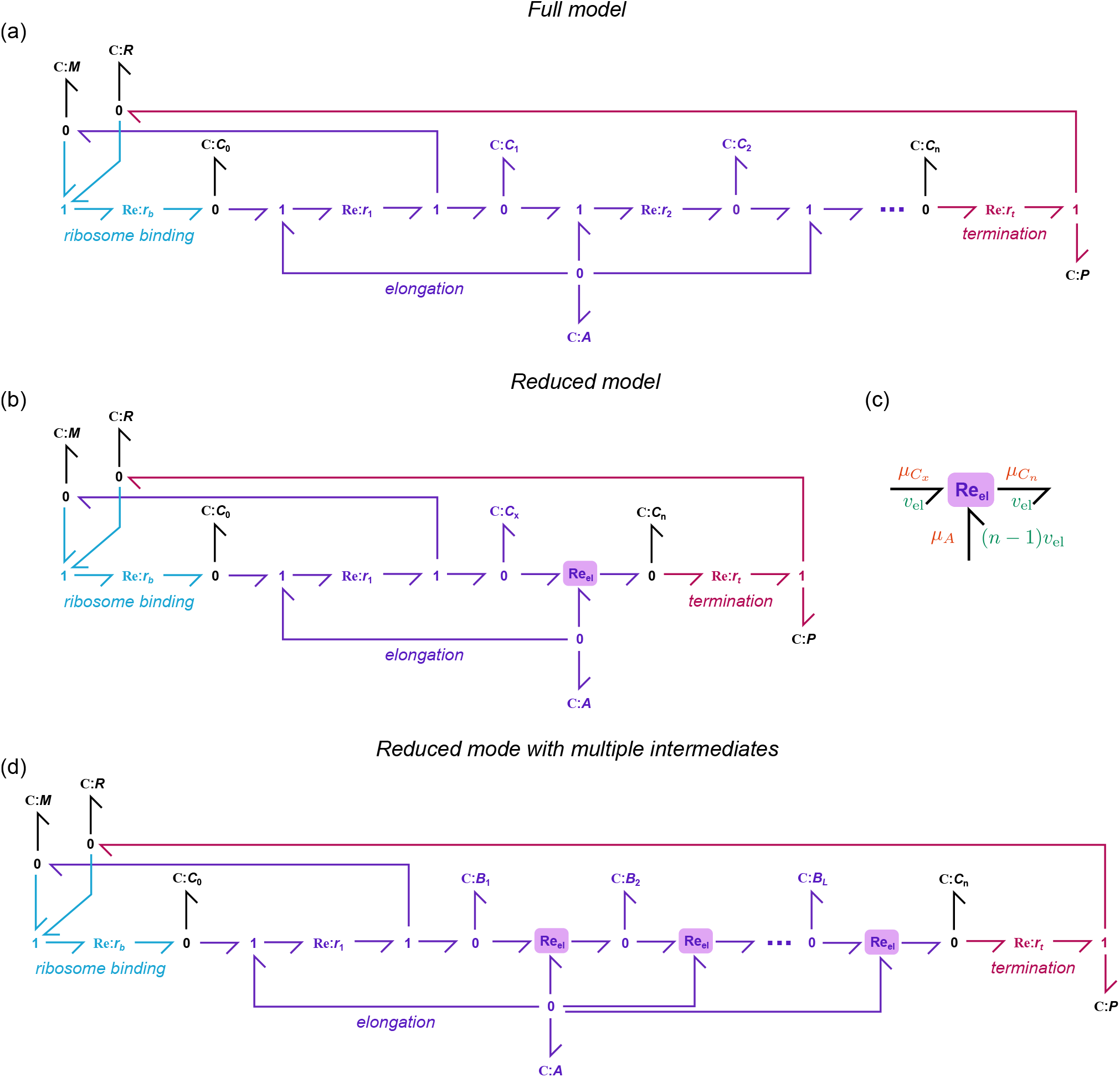
Bond graph models of translation. (a) Bond graph of the full translation model (§ 2.4.1); (b) bond graph of the reduced model (§ 2.4.2); (c) the Re_el_ component used to model elongation in (b); (d) bond graph of a reduced model with multiple intermediates.

#### 2.4.2 Model reduction

Since the number of ATP molecules per protein *n* can be on the order of thousands, it is computationally expensive to use the full model of translation in models of complex gene regulatory networks. Here, we outline a method for simplifying the full model while retaining its essential features. We group the complexes *C*_1_ to *C*_*n−*1_ into a lumped complex *C*_*x*_, giving rise to the reaction scheme

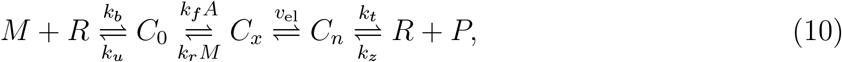

where *v*_el_ is a non-mass-action reaction representing the elongation rate. A bond graph of the system is shown in Figure 4b, where the elongation reaction is represented as a three-port reaction component (Figure 4c). In § A.2.1 of the electronic supplementary material, we show that under conditions where the dynamics of *C*_1_ and *C*_*n*_ are slower than those of the complexes in between, a quasi-steady-state approximation can be applied and the elongation rate is

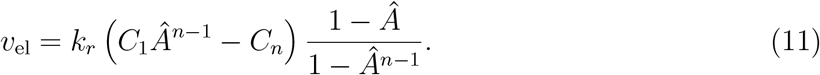

We show in § A.2.2 of the electronic supplementary material that this equation can be represented as a three-port bond graph component (Figure 4c) with the rate law

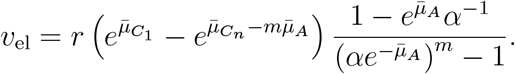

The lumping of intermediate complexes also requires the scaling of parameters and initial conditions, discussed further in § A.2.3 of the electronic supplementary material. In cases where the dynamics need to be better matched, it is also possible group the complexes into multiple lumps (Figure 4d; see § A.2.4 in the electronic supplementary material for details).

### 2.5 Gene expression and regulation

We construct a model of gene expression and regulation by coupling together the earlier models of transcription and translation (Figure 5). Since bond graphs are graphical by nature, they enable a modular approach to modelling biological systems. Using this approach, models of subsystems – in this case the transcription and translation modules – can be re-used in larger models without the need to manually re-derive the equations. Here, we couple bond graph modules using the approach in Gawthrop and Pan [24], where **C** components corresponding to shared species are exposed as ports within the transcription and translation modules. Degradation processes for mRNA and proteins are added to avoid overaccumulation (Figure 5c); the details are discussed in § A.3 of the electronic supplementary material. The full model equations are given in § A.4 of the electronic supplementary material.

**Figure 5:**
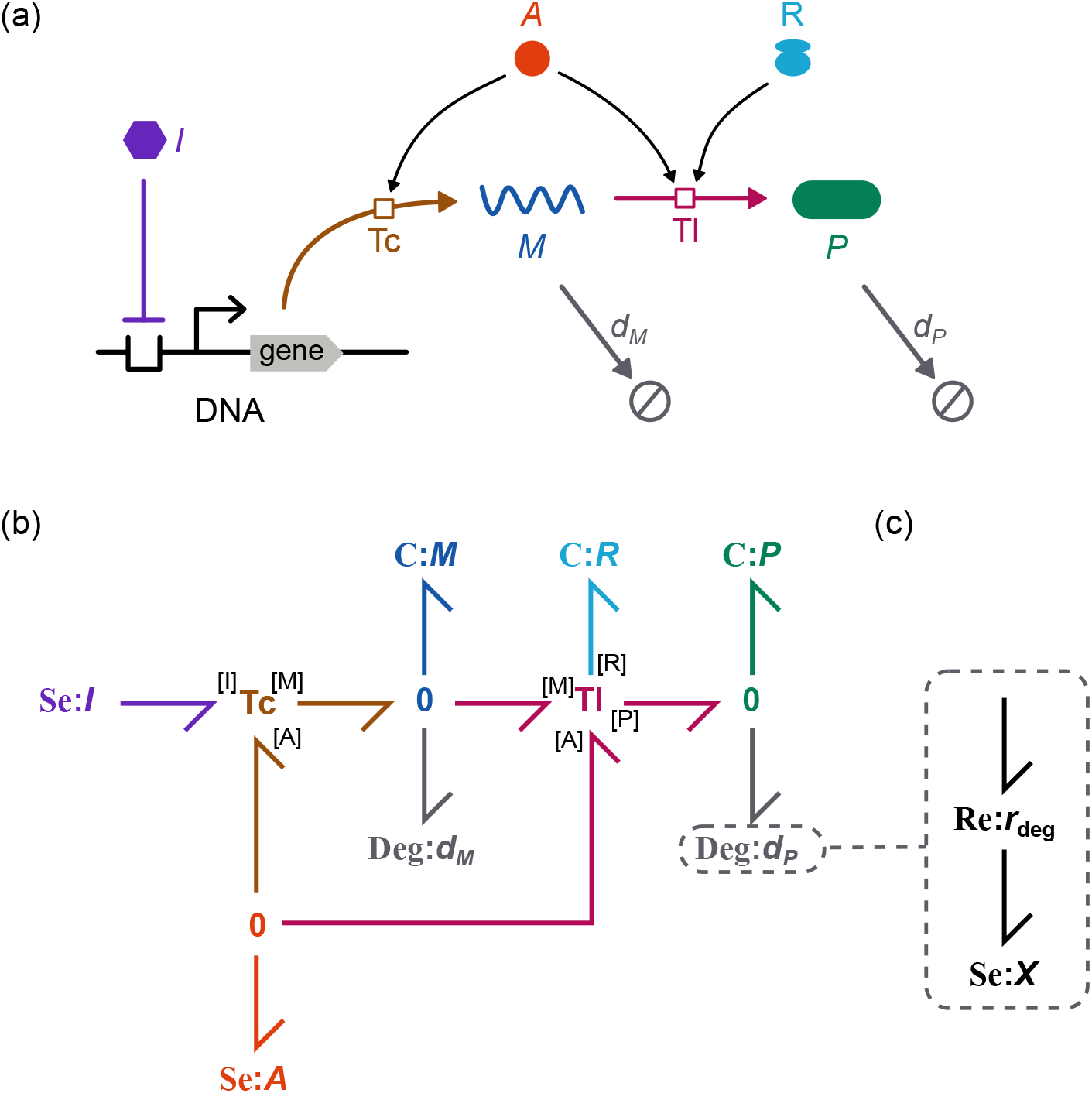
Model of gene expression. (a) Schematic and (b) bond graph model of gene expression. Degradation for mRNA and protein are modelled using the modules in (c). The transcription (Tc) module is shown in Figure 3 and the translation (Tl) module is shown in Figure 4b.

### 2.6 Synthetic biology motifs

Bond graphs of these modules are shown in Figure 6. These consist of an instance of the gene expression module **GE** for each gene, but with additional connections between the genes. In particular, the protein of one gene expression system acts as the repressor for another gene expression system. These connections are shown by the coloured bonds in Figure 6. Also, since all genes require energy and ribosomes to be expressed, these resources must be shared between modules. This is modelled by connecting the modules to **0** junctions (grey bonds in Figure 6) to ensure mass conservation.

**Figure 6:**
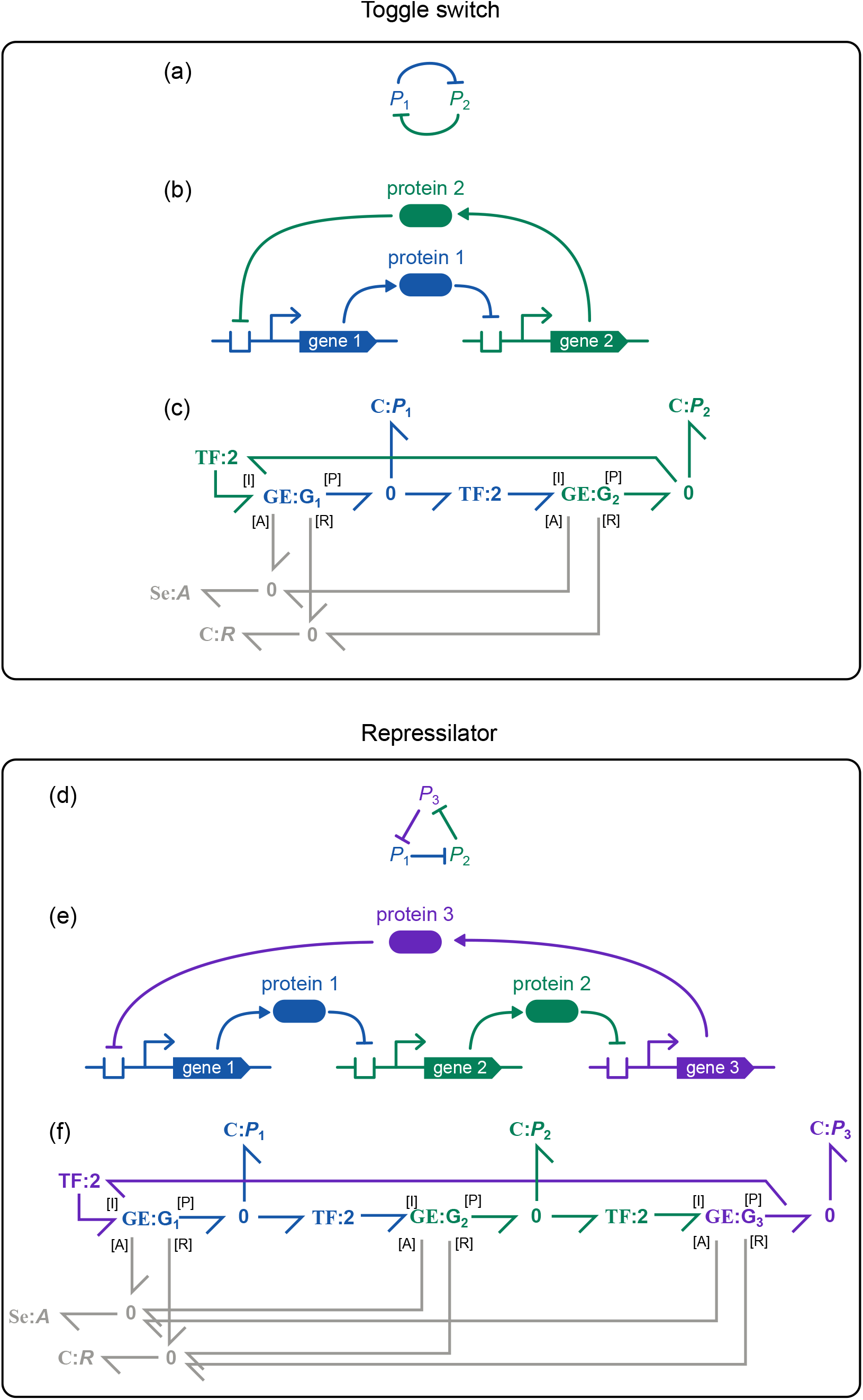
Models of synthetic biology motifs. The bond graph model of (a–c) the toggle switch; and (d–f) repressilator. For each panel, the relationships between proteins are shown on top (a,d), the mechanisms of interaction are displayed in the middle (b,e) and bond graphs are shown on the bottom (c,f). The **GE** modules refer to the model in Figure 5b.

Gene regulatory networks often require binding cooperativity to enhance their sensitivity to inputs [36]. This is incorporated by using a **TF** component (§ 2.2.4) with modulus *h* to model the binding of multiple inhibitors to prevent transcription, resulting in a transcription rate of

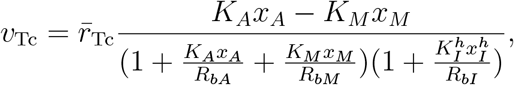

A Hill coefficient of *h* = 2 is used, consistent with previous models [4, 20].

## 3 Results

### 3.1 Comparison of full and reduced models

#### 3.1.1 Elongation

To ensure the validity of the model reduction scheme in § 2.4.2, we use dynamic simulations to compare the output of the full and reduced models of translation. We first examine the elongation reactions in isolation to verify that the simple model matches the elongation rate of the full model at steady state. Figure 7a shows the results obtained from simulating the full model of elongation (Eq. (S3) of the electronic supplementary material), compared to the analytical form of the elongation rate (Eq. (11)). For simplicity, we use a chain length of *m* = 5 for these simulations, but the results generalise to longer chain lengths (see below). As expected, the full and reduced models match very well. The steady-state amounts of the intermediate complexes *X*_1_, *X*_2_, *X*_3_ and *X*_4_ are plotted in Figure 7b, and also match closely. We next investigate whether the model reduction scheme holds for different chain lengths *m* (Figure 7c). The elongation rate decreases, and ultimately plateaus at high *m*; and the reduced model is able to predict this behaviour. Thus, the reduced model is able to accurately match the steady-state behaviour of the full model under a wide range of conditions.

**Figure 7:**
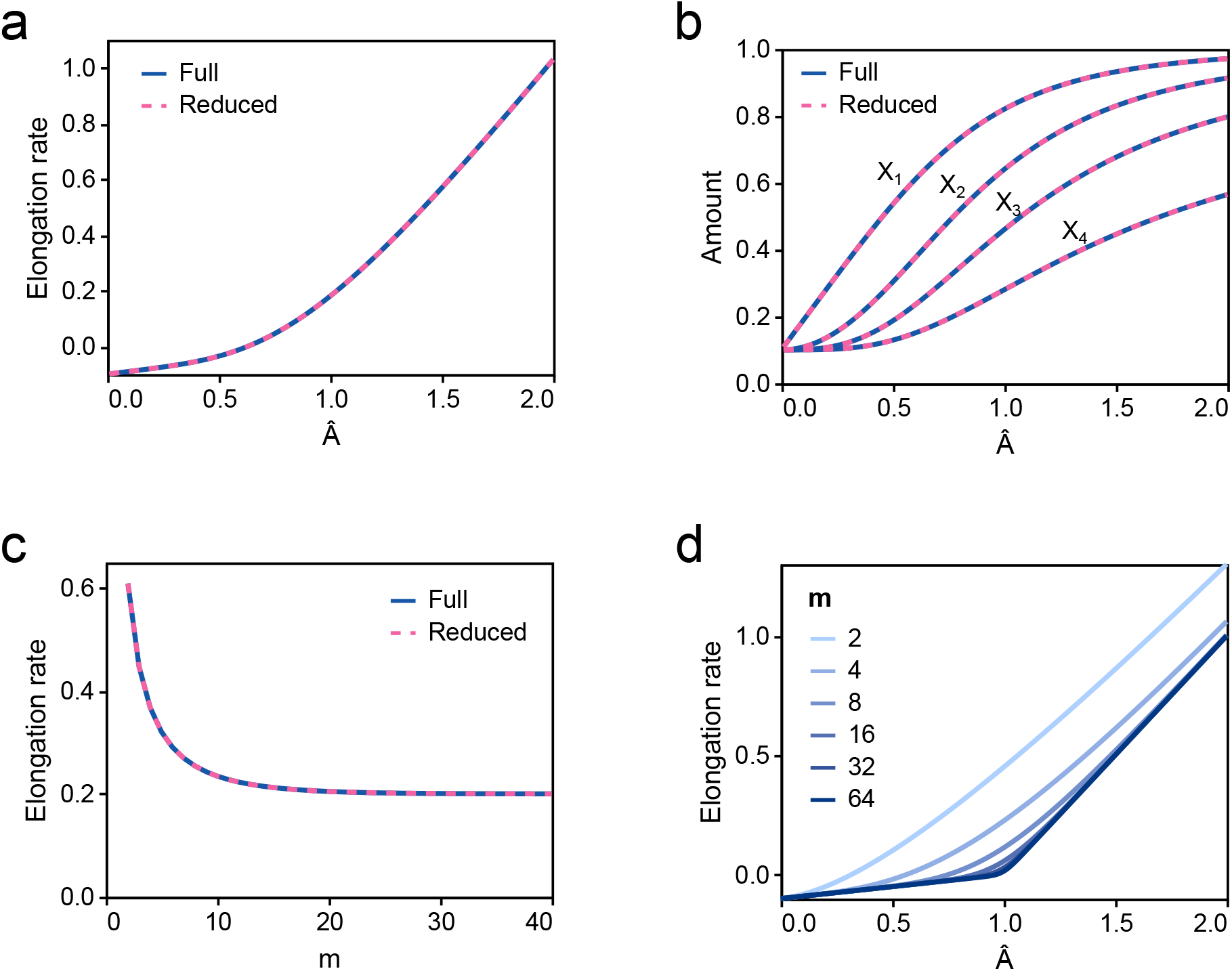
A simplified model of elongation. Steady-state comparisons for (a) elongation rates and (b) complex amounts are plotted for different values of Â. (c) Steady-state comparison of elongation rates for different values of *m*. The simulations in (a)–(c) use the kinetic parameters *k*_*f*_ = 1, *k*_*r*_ = 1, *C*_0_ = 1.0 and *C*_*n*_ = 0.1, with *A* set to 1.2 for (c). (d) A plot of the analytical solution for different values of *m*. Models are run for *t* = 50 minutes to achieve steady state.

Figure 7d shows the relationship between elongation rate (Eq. (11)) and *Â* for different chain lengths *m*. For shorter chain lengths, the elongation rate is smooth with respect to *Â*. However, as predicted by equation Eq. (S7) of the electronic supplementary material, for large *m*, the curve is almost piecewise linear with different slopes on either side of *Â* = 1.

#### 3.1.2 Translation

While the above results confirm similarities between the full and reduced models at steady state, we are interested in comparing their behaviours when included in a dynamic model. Thus, we next investigate whether the dynamics of translation are similar between the full and reduced models. To simulate the models under biological conditions, the models are parameterised based on previous experimental and modelling work (see Appendix B in the electronic supplementary material). We simulate the dynamics of the full (Figure 4a) and reduced models (Figure 4b), and compare them for different energy levels and protein lengths (Figure 8). Models are initialised from an inactivated state with very small amounts of ribosome-mRNA complexes, and the evolution to an activated state is compared between models. We test the match between models for a range of energetic states *A*, with *A* = 5.8 *×* 10^6^ molecules/cell corresponding most closely to an energy-rich environment (§ B.1 in the electronic supplementary material). We also test the quality of match for different chain lengths *n*.

**Figure 8:**
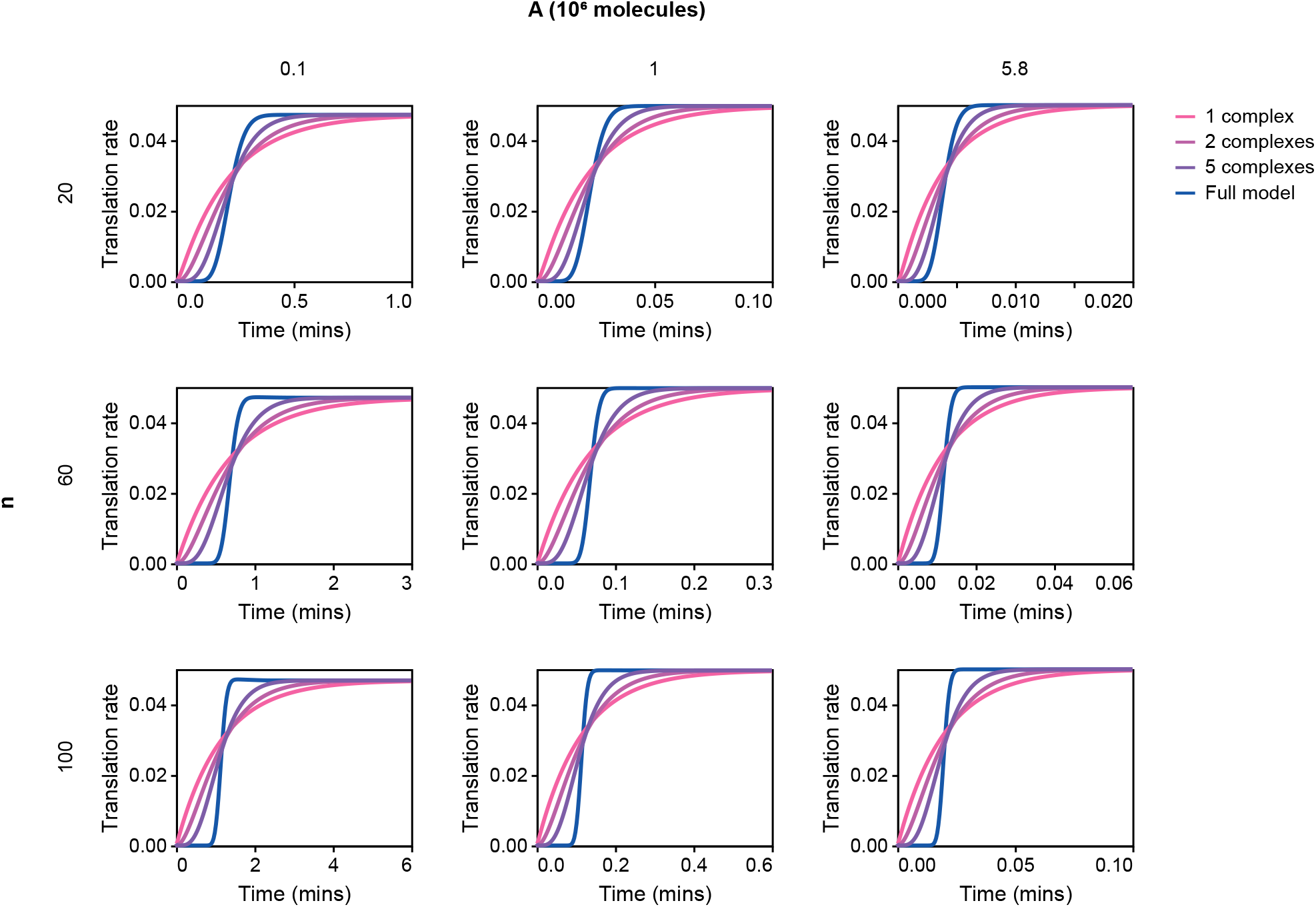
Comparison of the dynamics of translation rate. The energetic state *A* is varied horizontally, and the chain length *n* is varied vertically. The translation rate is given in proteins per minute. Simulations are run with 10 mRNA molecules, 5000 ribosomes and 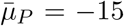 to approximate the immediate removal of proteins.

In all cases, the reduced model approaches the same translation rate as the full model. However, the models exhibit different dynamics. Whereas the full model has a more step-like response to a sudden increase in mRNA, the reduced model approaches the steady-state translation rate more gradually. As one would expect, the match between dynamics becomes poorer for larger values of *n*. Nonetheless, both the full and reduced models approach their steady states at similar timescales for a wide range of chain lengths. Due to the similarities in steady-state behaviour and timescales, we believe that the simple model is an adequate approximation, and we indeed show in § 3.2 that the differences in dynamics result in very minor differences at the level of gene expression. In circumstances where better matching with the dynamic behaviour is required, one could lump elongation into multiple intermediate states rather than a single complex. This approach is analogous to epidemiological models with multiple exposed or infectious compartments [37] and models of multi-stage transport processes [38]. Simulations of two and five lumped complexes are displayed in Figure 8, and show a gradual improvement of fit with the number of intermediates.

We also find that the reduced model is computationally efficient compared to the full model. A benchmark of runtimes is presented in Figure 9. As expected, the full model takes longer to run as the number of state variables increases; the approximately linear gradient at high *n* indicates a rapidly increasing runtime with protein length. In contrast, the time taken to run the reduced model remains stable even for higher protein lengths. At the longest tested protein length of *n* = 400, a 300 *×* speedup is observed for the reduced model, relative to the full model. We expect even greater speedup for realistic protein lengths of over 1000 ATP molecules per protein.

**Figure 9:**
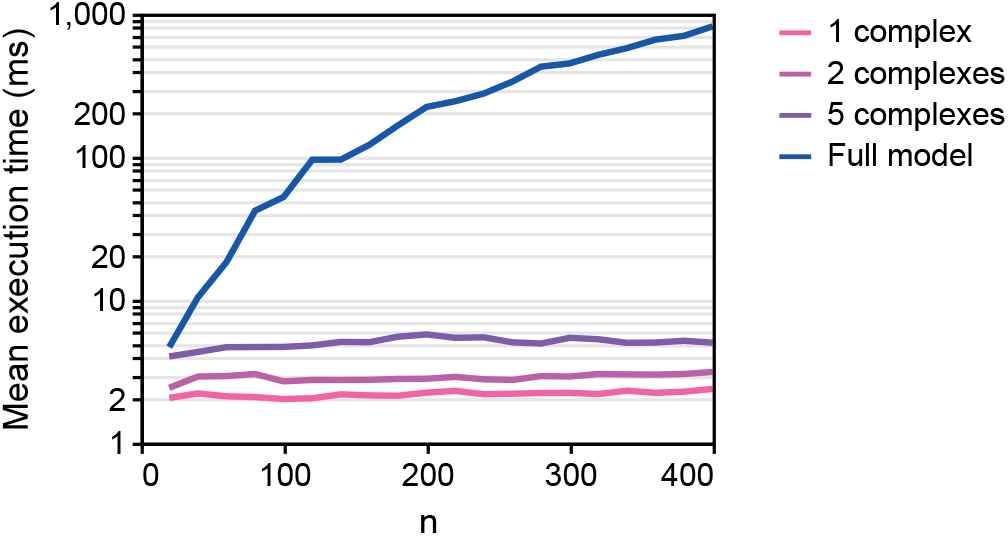
Benchmark of full and reduced models. Simulations are run on the translation model and the mean execution time is plotted for different protein lengths *n*. Models were solved using the CVODE Backward Differentiation Formula method [39] with a relative tolerance of 10^*−*9^ and absolute tolerance of 10^*−*12^. Models were repeatedly simulated over either 1000 samples, or until 1 minute had elapsed in real time. The code was simulated on a Windows 10 server with 4 CPU cores (AMD EPYC 7702 processor).

In Appendix D of the electronic supplementary material, we verify that the bond graph model is thermodynamically consistent and analyse the energetics of gene expression under different energy availabilities.

#### 3.2 Gene expression

Models of gene expression are constructed by coupling models of transcription and translation (Figure 5). We simulate the induction of a gene, where an inhibitor is initially present at high concentrations before being removed (Figure 10a). As expected, following the removal of the inhibitor, mRNA levels rise followed by a rise in the the protein. The counts of these molecules are within the ranges observed experimentally [40]. In § C.1 of the electronic supplementary material, we show that under the assumption of excess energy, the steady-state mRNA and protein numbers are

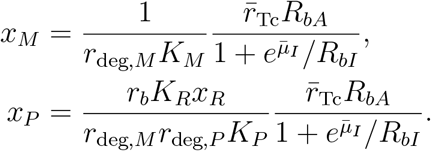

**Figure 10:**
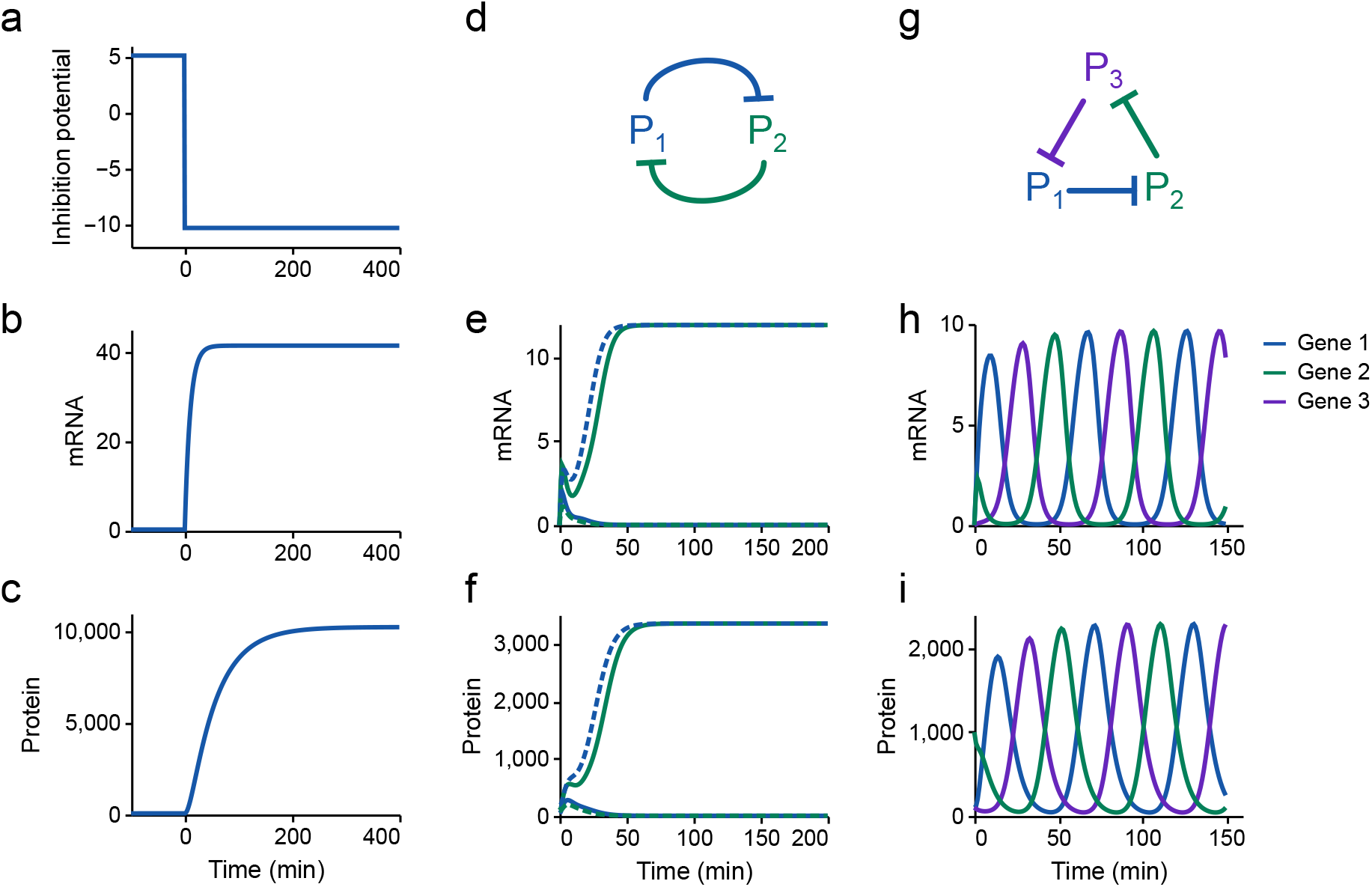
Simulations of synthetic circuits. (a–c) Simulations of a single gene. The inhibition potential *μ*_*I*_ *−* ln(*R*_*bI*_) is reduced from a high to low value at *t* = 0, indicating a switch from an inactive to active state. (d–f) Simulations of the toggle switch. Two simulations (represented using solid and dashed lines) are run with different initial conditions. The steady states are dependent on the initial condition, indicating bistability. (g–i) Simulations of the repressilator circuit.

With the parameters in the model, *x*_*M*_ *≈*41 and *x*_*P*_ *≈*10351, which is in good agreement with Figure 10b–c. The power dissipation rate is plotted against time in Figure S3 of the electronic supplementary material, which indicates that protein synthesis and degradation account for the majority of energy dissipated within the circuit.

To ensure that the model reduction scheme in § 2.4.2 is valid, the same models are simulated with the full model of translation (Figure 4a) in place of the reduced models. The resulting mRNA and protein levels are virtually identical to that of Figure 10 (Figure 11), and the quality of match is similar for the models of the toggle switch and repressilator. Thus, we expect that translation operates near steady state under biological conditions, and that the loss of dynamics from model reduction is unimportant under these conditions.

**Figure 11:**
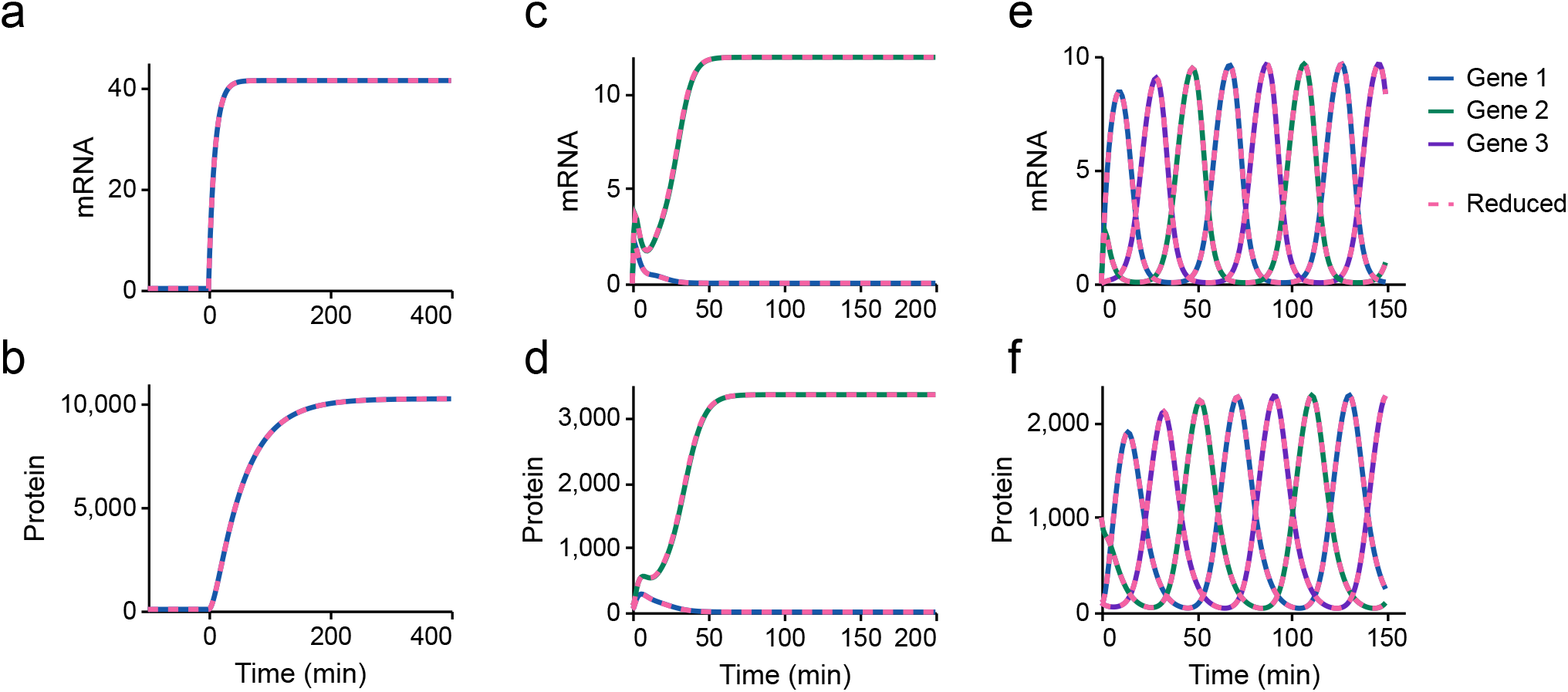
Comparison of full and reduced models of synthetic circuits. Models of synthetic circuits using the full translation model (solid lines) are compared to the corresponding models using the reduced translation model (dashed lines). (a–b) Induction of a single gene; (c–d) toggle switch; (e–f) repressilator.

In Figure 12, the translation rate is plotted against the amount of energy and ribosomes.

**Figure 12:**
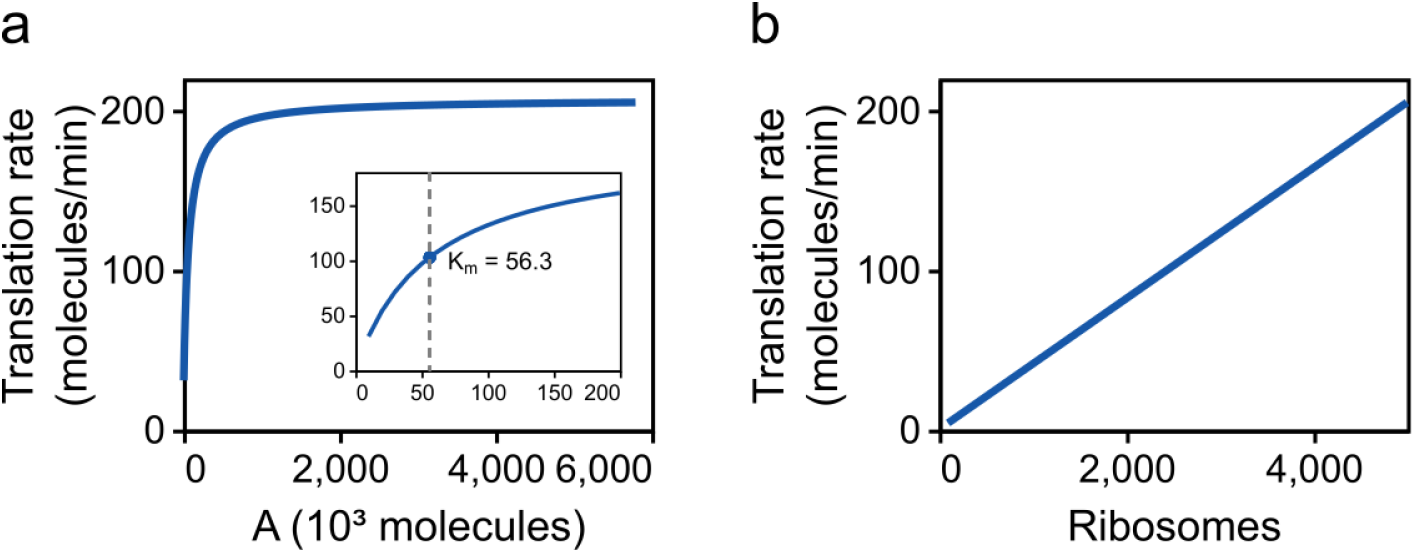
The effect of resource abundance on translation rate. The translation rate is plotted against (a) amount of energy; and (b) the number of ribosomes. The inset in (a) shows the translation rate for low amounts of energy.

Decreasing the amount of energy has little effect on the translation rate (Figure 12a) until the amount of energy is very low, reflecting energy starvation conditions. The translation rate reaches 50% of its maximum value at *A* = 5.63 *×* 10^4^ molecules (Figure 12a; inset), which corresponds to a concentration of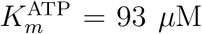. It is reassuring that this value is on the same order of magnitude as experimental measurements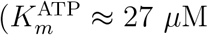 ; Jewett et al. [41]) despite the fact that the model has not been fitted to such data. Decreasing the number of ribosomes reduces the translation rate linearly (Figure 12b). This appears to be consistent with studies that hypothesise ribosomes are a limiting factor for translation [42]. We also show in § C.1 of the electronic supplementary material that the steady-state translation rate is linearly dependent on ribosome number at high energy levels.

#### 3.1 Resource dependence of synthetic circuits

##### 3.3.1 Toggle switch

We next explore the behaviours of synthetic biology circuits when modelled with thermodynamic dependencies in mind. A model of the toggle switch (Figure 6a–c) is constructed by coupling together two mutually inhibiting genes. We perform simulations of the toggle switch under two different initial conditions (Figure 10d–f), corresponding to the activation of protein 1 (dashed lines) and protein 2 (solid lines). As expected, the model reproduces the multistability of the switch observed experimentally [3].

As discussed in Section 1, synthetic circuits do not operate in isolation, and are affected by background processes such as the allocation of energy and ribosomes. We examine the effects of limiting either energy (*A*) or ribosome (*R*) availability on the performance of the toggle switch. The performance of the toggle switch is characterised using two measures:

1. the *bistability index*, defined as the steady-state amount of *P*_1_ in the active steady-state relative to its amount in the inactive state [43].
2. the *switching time*, defined as the time taken for both proteins to reach their steady state [3]. To quantify the switching time, we define the distance from the steady state as

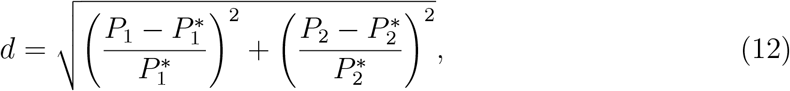

where 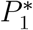 and 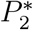 are the steady-state amounts of proteins 1 and 2 respectively. The switching time is the time taken for the model to reach a distance of *d <* 0.01.

Figure 13 shows the effect of energy and ribosomes on the behaviour of the toggle switch. We find that ribosomes are the primary determinant of toggle switch bistability, as the bistability index varies by approximately 30-fold over the range of ribosome amounts considered. The steady states of the toggle switch under energy-rich conditions are analysed in § C.2 of the electronic supplementary material, verifying this result. In contrast, both the energy and ribosomes affect the switching time. A higher energy availability reduces the switching time as expected, since energy accelerates the dynamics of translation. Surprisingly, the number of ribosomes has a nonmonotonic effect on switching time, and achieves a minimum value between 3000–6000 ribosomes. This leads to potential trade-offs in performance at high numbers of ribosomes; increasing ribosomal availability increases the bistability index, but also increases the switching time.

**Figure 13:**
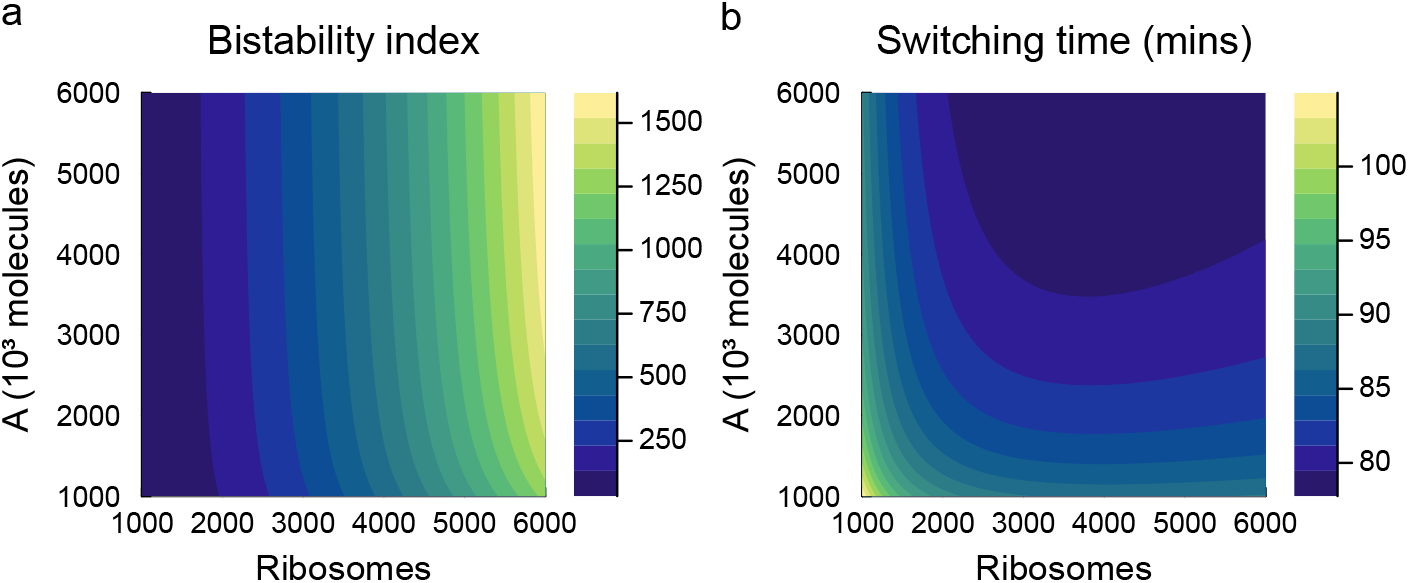
Simulations of the toggle switch under varying energy and ribosome levels. Simulations are run with an initially high amount of protein for gene 1, resulting in a steady state favouring this gene. (a) The bistability index, defined as the steady-state amount of protein 1 in its active state relative to its inactive state; (b) switching time, defined as the time required for the system to be within 1% of its steady-state value (Eq. (12)).

#### 3.3.2 Repressilator

We next study the dynamic behaviour of the repressilator. Simulations of the repressilator show oscillations between the three inhibitory proteins, in line with the expected function of the circuit (Figure 10g–i). In Figure 14, we explore the behaviour of the circuit under different energy and ribosome availabilities, characterising the amplitude and period of the steady-state oscillations. The amplitude of oscillations is primarily determined by the number of ribosomes; a reduction of ribosomes from 5000 to 1000 results in an approximately 7-fold decrease in oscillation amplitude (Figure 14a). Both energy and ribosome availability contribute to the period of oscillations. Increasing energy availability accelerates the dynamics of translation, increasing the frequency of oscillations, whereas increasing the number of ribosomes decreases the frequency of oscillations. Interestingly, similarities can be drawn with results from the toggle switch, where measures associated with the amplitude of protein changes are mostly impacted by ribosome availability, whereas temporal behaviours are determined by both energy and ribosome availability.

**Figure 14:**
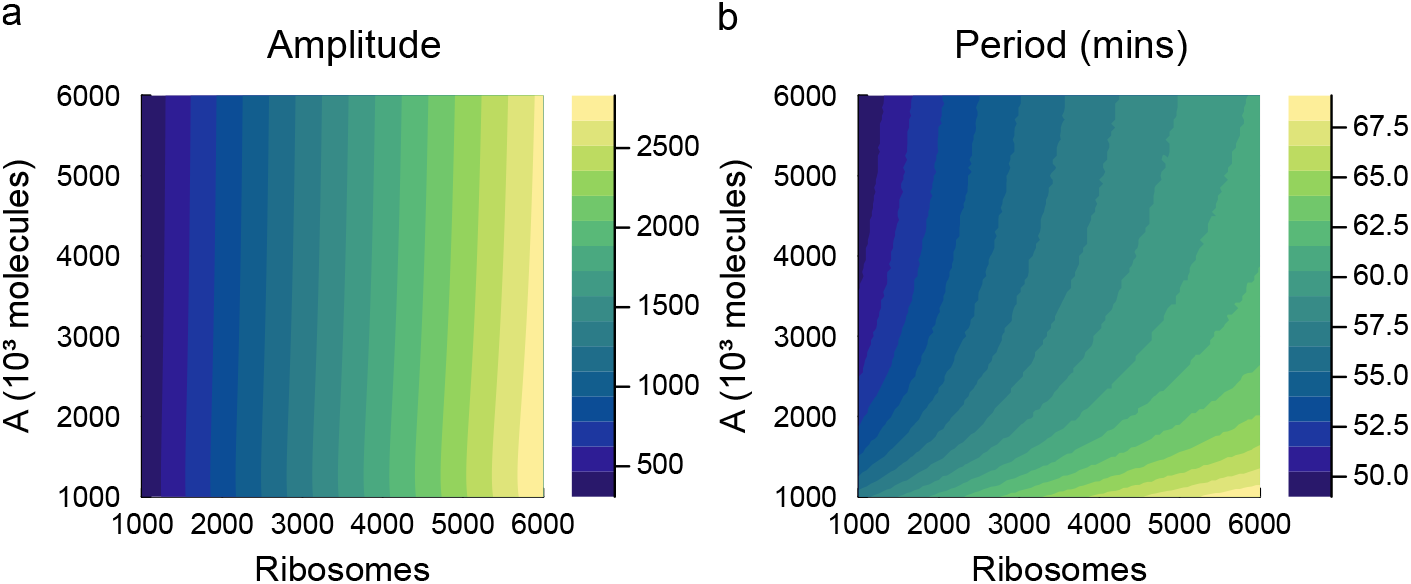
Simulations of the repressilator under varying energy and ribosome levels. Simulations are run until a limit cycle is reached, and the (a) amplitude and (b) period of oscillations are recorded.

### 3.4 Parameter heterogeneity

A characteristic of cell populations is that they may experience different local environments and therefore display heterogeneity in their behaviour [44]. Because our simple model of gene expression is computationally efficient, it is well-suited to exploring the effects of heterogeneity within cell populations. Accordingly, we explore the effects of parameter heterogeneity on the behaviour of the repressilator (Figure 6d–f). Note that these investigations are intended to be a proof-of-concept for simulating cell populations since they assume cells operate independently of each other. In reality, cells influence one another through the transmission of extracellular signals and the competition for nutrients and resources.

We first investigate the impact of heterogeneity in energy availability on repressilator behaviour. Nakatani et al. [45] find that ATP concentrations vary substantially in populations of *E. coli*. To simulate the effect of this heterogeneity, we fit a normal distribution (mean = 3.76 mM, std = 1.64 mM) to Fig. 1A of Nakatani et al. [45]. These concentrations are converted to a number of molecules, assuming a cell volume of 1 *μ*m^3^ [20]. We obtain samples of energy levels using a truncated normal distribution with a minimum of *A* = 10^4^ molecules per cell. The resulting behaviour for 10000 samples is summarised in Figure 15a–d. Due to the relative insensitivity of the repressilator model to energy levels, the effect on the amplitude of oscillations was small, with the vast majority of samples having amplitudes between 2200–2400 (Figure 15b). The effect on the period of oscillations was similarly small, with most samples having a period of between 60–66 mins (Figure 15c).

**Figure 15:**
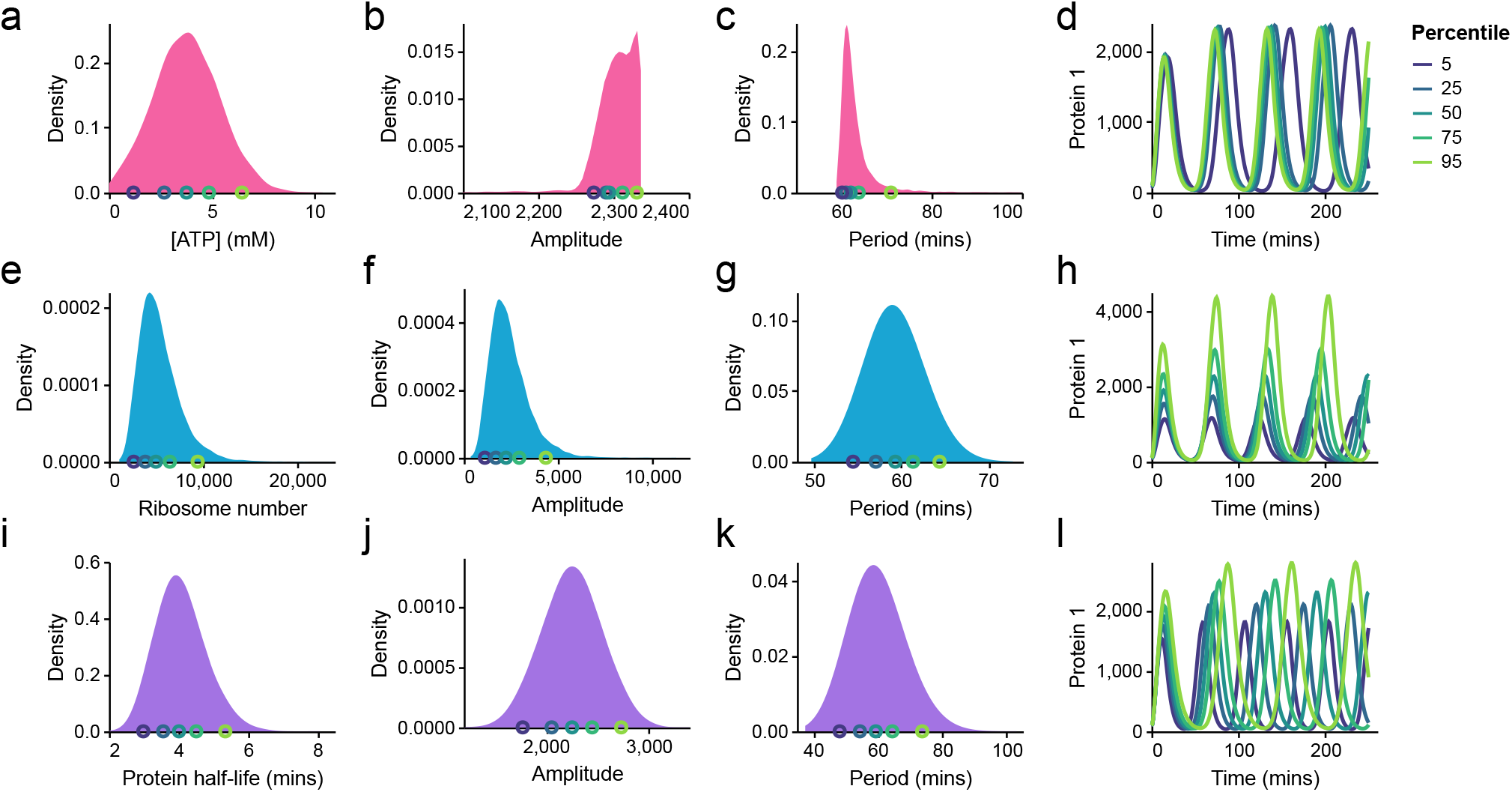
Effects of parameter heterogeneity on repressilator function. Each row corresponds to distributions extracted from running 10000 simulations, with a parameter varied by sampling from a distribution. (a–d) Effects of heterogeneity in ATP concentration. Changes in ATP concentration were modelled by adjusting the parameter *μ*_*A*_ (14 samples were excluded from analysis as they resulted in no oscillations due to insufficient energy levels). (e–h) Effects of heterogeneity of ribosome amount, which are modelled by changing the initial ribosome number. (i–l) Effects of heterogeneity in protein decay rates, incorporated by adjusting the parameter *r*_deg,*P*_. For each row, simulations corresponding to the 5th, 25th, 50th, 75th and 95th percentiles of the parameters are plotted on the right (d,h,l). The colours of the timecourses correspond to the circles of the same colour in the distributions.

Next, we consider heterogeneity in ribosome amounts. We assume that the number of free ribosomes per cell follows a similar distribution to protein counts in cells. Milo and Phillips [40] estimated that protein counts have a coefficient of variation of approximately 0.4 in bacteria. Thus, we sample ribosome counts from a log-normal distribution around a median free ribosome count of 5000 (Figure 15e). This heterogeneity has a marked impact on the amplitudes of oscillation (*∼*8-fold variation; Figure 15f), and a mild impact on the period (*∼* 25% variation; Figure 15g).

We finally explore the effect of variations in protein degradation rates. Since the protein half-life has been shown to vary 2–3 fold among populations of *E. coli* [46], we assume the protein half-life is log-normally distributed with a median half-life of 4 minutes and a 95% confidence interval spanning from 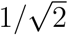 to 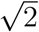 times the median. We also assume that protein half-lives are perfectly correlated between proteins, caused by global fluctuations in cell state [44]. Using these proof-of-concept simulations, heterogeneity in protein degradation results in a wide range of variation in both the amplitudes and periods of oscillations, with approximately 2-fold variation in their values (Figure 15i–l).

## 4 Discussion

In this study, we develop a new bond graph model of gene expression. In contrast to previous models in the literature [17, 18, 20, 21], our model is both simple and thermodynamically consistent, making it suitable for understanding and designing synthetic circuits in cell populations. By coupling together multiple instances of our gene expression module, we are able to reproduce behaviours of two foundational synthetic biological circuits: the toggle switch and repressilator [3, 4]. However, the approach is generalisable to other synthetic circuits involving protein synthesis, such as antithetic feedback [47] and could be used to study the effects of energy burden in such control systems. Our approach is based on bond graphs, a general framework for modelling biophysical systems. Due to their physical consistency, commonly neglected phenomena such as resource competition and retroactivity are automatically built into bond graph models [8, 10]. Additionally, their modular and hierarchical structure allows modellers to systematically couple models of synthetic circuits into larger circuits [28]. Thus, we believe that bond graphs hold great potential in aiding the design of circuits in synthetic biology.

Our model of gene expression is based on a model reduction scheme to simplify the elongation reactions in translation (§ 2.4.2). It is common for other simple models of translation to assign rate laws based on empirical fits to data [4, 17]. By contrast, our model reduction approach is biophysically motivated, based on a detailed reaction network containing molecules and processes known to be involved in translation. As a result, our model reduction scheme preserves important constraints such as mass and energy conservation. We show that the reduced model behaves similarly to the full model at steady state and reaches the steady state on similar timescales (Figures 7–8). Furthermore, we find that the loss of dynamics in translation is unimportant when simulating synthetic gene circuits (Figure 11). The reduced model is also computationally efficient, able to achieve runtimes hundreds of times (and possibly up to thousands of times) faster than the full model (Figure 9). This speedup is quite significant when scaling up to large gene regulatory networks or agent-based models of large cell populations. Given that the addition of monomers to macromolecules is prevalent in biology, we believe that our model reduction scheme would also be applicable to processes such as transcription, glycogenesis and fatty acid polymerisation [48].

In spite of the constraints that energy places on cells, we still have an incomplete picture of how energy constrains cellular processes and how the cell distributes its energy budget between processes [49]. While previous works have studied the effects of ribosome limitation on the function of synthetic circuits [50, 51], to the best of our knowledge, our model is the first to incorporate the known thermodynamics of ATP hydrolysis into a lumped parameter dynamic model of translation. Our model predicts a relative insensitivity of gene expression to energy, which appears to be supported by experimental measurements [41]. Our model is also able to analyse the thermodynamic costs associated with translation. We find that the majority of energy supplied through ATP hydrolysis is dissipated during elongation rather than being stored in the peptide bonds in proteins (Appendix D in the electronic supplementary material). This inefficiency has previously been attributed to proofreading mechanisms [52]; more detailed modelling of these mechanisms is required to understand performance-cost trade-offs in translation.

Our model also makes predictions about how energy and ribosome availability affect the behaviour of synthetic circuits. Interestingly, the number of free ribosomes is the primary determinant of measures associated with protein counts, whereas both energy and ribosome availability contribute to the dynamic behaviour of the circuits. We also explore the potential effects of parameter heterogeneity in cell populations. Informed by empirical measurements, we find that variations in energy levels have a limited impact on the behaviour of the repressilator (Figure 15). However, variability in both ribosome availability and protein degradation rates result in high degrees of variation in both the amplitude and period of oscillations. Furthermore, each parameter influences different aspects of the function of the repressilator. While ribosome availability greatly influences the amplitude of oscillations, the protein degradation rate has a greater impact on the frequency of oscillations. Thus, the parameter variability that exists between cells may be responsible for cell-to-cell variability in the performance of synthetic circuits, in the absence of mechanisms of intercellular interaction [3, 53].

While we explore parameter heterogeneity as a proof-of-concept for modelling cell populations, mathematical questions remain on how to incorporate mechanisms of cell-to-cell communication. Future work will focus on incorporating our model of gene expression into an agent-based model. While the model in this study is broadly applicable to synthetic biology, the bioluminescence pathway in bacteria is a promising model system to engineer for several reasons, including (i) the ease of measuring light as the output, (ii) the involvement of quorum sensing in coordinating population-level behaviours and (iii) the use of two competing pathways for its induction [54]. Furthermore, the multi-physical nature of bond graphs allows them to naturally account for the conversion of chemical energy to light [24]. Modelling the intracellular regulatory mechanisms associated with quorum sensing alongside intercellular communication has the potential to enable the engineering of synchronised behaviour of cell populations [15, 55].

Due to the simple nature of the model, certain details of gene regulation have been omitted. While we assume that elongation involves a single reaction per ATP molecule, tRNA is in reality charged by amino acids through elongation factors, which involves a more complex series of reactions [56]. The elongation rate is dependent on tRNA availability, and therefore specific to each codon. We also lump energy into a single molecule, but in reality, it is composed of the hydrolysis of ATP into ADP, Pi and H^+^. Accounting for such details will be the subject of future work.

## Supporting information

Supplementary methods and results

## Acknowledgments

S.T.J. is supported by the Australian Research Council (Project No. DE200100988). P.J.G. would like to thank the Faculty of Engineering and Information Technology, University of Melbourne, for its support via a Professorial Fellowship. M.P. was supported by a Postdoctoral Research Fellowship from the School of Mathematics and Statistics at the University of Melbourne.

## Data availability

The bond graph models were simulated in Julia using the BondGraphs.jl package [57]. The code is available on GitHub at https://github.com/mic-pan/bg_gene_expression.

