## Supplementary methods and results for "Thermodynamically-consistent, reduced models of gene regulatory networks"

*Supplementary Material*

Michael Pan, Peter J. Gawthrop, Matthew Faria, Stuart T. Johnston

---

**D Energetic analysis** **11**

**E Supplementary figures** **13**

### A Model details

#### A.1 Transcription

The elementary reaction network for transcription is

$$r_1 : A + E \rightleftharpoons EA \tag{S1a}$$
$$r_2 : EA \rightleftharpoons E + M \tag{S1b}$$
$$r_3 : E + I \rightleftharpoons EI \tag{S1c}$$
$$r_4 : EA + I \rightleftharpoons EAI \tag{S1d}$$
$$r_5 : EI + A \rightleftharpoons EAI \tag{S1e}$$

where $E$ is the free enzyme and $EA$, $EI$ and $EAI$ are enzyme states. Under the assumption that reactions (S1c)–(S1d) are at quasi-equilibrium and have the same dissociation constant $K_{d,I}$, this represents non-competitive inhibition [1].

We apply the quasi-steady-state approximation [2], and therefore, the reaction network simplifies to

$$A \rightleftharpoons M$$

with the net reaction rate defined as

$$v = \bar{r}\frac{e^{\bar{\mu}_A} - e^{\bar{\mu}_M}}{(1 + \frac{e^{\bar{\mu}_A}}{R_{bA}} + \frac{e^{\bar{\mu}_M}}{R_{bM}})(1 + \frac{e^{\bar{\mu}_I}}{R_{bI}})}, \tag{S2}$$

where

$$\bar{r} = \frac{r_1 r_2}{r_1 + r_2},$$

is the rate constant and

$$R_{bA} = \frac{(r_1 + r_2)K_{EA}}{r_1 K_E},$$
$$R_{bM} = \frac{(r_1 + r_2)K_{EA}}{r_2 K_E},$$
$$R_{bI} = K_I K_{d,I},$$

are the binding constants. Since the grouped parameters $\bar{r}$, $R_{bA}$, $R_{bM}$ and $R_{bI}$ are sufficient to define Eq. (S2), we treat these as the parameters for transcription.

#### A.2 Translation

##### *A.2.1 Model reduction scheme*

To simplify elongation, we consider the reactions associated with elongation after the first step:

$$C_1 \underset{k_r}{\overset{k_f A}{\rightleftharpoons}} C_2 \underset{k_r}{\overset{k_f A}{\rightleftharpoons}} \dots \underset{k_r}{\overset{k_f A}{\rightleftharpoons}} C_n. \tag{S3}$$

To establish a connection to the reactions involved in translation, we make the simplifying assumption that $C_1$ and $C_n$ are boundary species whose concentrations change slowly relative to the other complexes in the chain. We expect this assumption to be reasonable since binding dynamics are generally slower than elongation dynamics. Nonetheless, we outline a method in § A.2.4 for achieving a better match in cases where this assumption does not hold. Following the quasi-steady-state approximation, $\mathrm{d}C_i/\mathrm{d}t = 0$ for $i \in \{2, \dots, n-1\}$. Then

$$\frac{\mathrm{d}C_i}{\mathrm{d}t} = k_f A C_{i-1} + k_r C_{i+1} - (k_f A + k_r) C_i = 0, \quad i \in 2, \dots, n-1. \tag{S4}$$

To solve this system of equations, we treat $C_i$ as a discrete sequence and analyse its $z$-transform. For simplicity, we define $x_i = C_{i+1}$ for $i = 0, \dots, n-1$ and $X(z)$ as the $z$-transform of $x_i$. The sequence follows the difference equation

$$k_r x_{i+2} - (k_f A + k_r) x_{i+1} + k_f A x_i = 0, \quad i \geq 0.$$

Taking the $z$-transform of both sides, we find

$$k_r z^2 X(z) - (k_f A + k_r) z X(z) + k_f A X(z) = p(z),$$

where $p(z)$ is a quadratic used to initialise the sequence. We then factorise the quadratic on the left-hand side:

$$k_r (z-1)(z-\hat{A}) X(z) = p(z),$$

where $\hat{A} = k_f A / k_r$. Using partial fraction decomposition,

$$X(z) = \frac{az}{z-1} + \frac{bz}{z-\hat{A}},$$

for some constants $a$ and $b$. Hence, by taking the inverse $z$-transform,

$$C_i = x_{i-1} = a + b\hat{A}^{i-1}, \quad i \geq 1. \tag{S5}$$

The constants $a$ and $b$ can be solved for substituting the boundary values $C_1$ and $C_n$:

$$a = -\frac{C_1 \hat{A}^{n-1} - C_n}{1 - \hat{A}^{n-1}},$$
$$b = \frac{C_1 - C_n}{1 - \hat{A}^{n-1}}.$$

Hence, the elongation rate is

$$v_{\mathrm{el}} = k_f A C_i - k_r C_{i+1} = k_r a (\hat{A} - 1) = k_r \left( C_1 \hat{A}^{n-1} - C_n \right) \frac{1 - \hat{A}}{1 - \hat{A}^{n-1}}. \tag{S6}$$

We make a few remarks on this expression:

1. In the case where the lumped complex contains a single species ($n = 2$), the elongation rate reduces to the law of mass action.
2. If $\hat{A} \ll 1$, the elongation rate is well approximated by the law of mass action:

$$v_{\mathrm{el}} \approx k_r \left( C_1 \hat{A}^{n-1} - C_n \right).$$

3. If $\hat{A} \gg 1$ and $C_n \ll \hat{A}^n C_1$, the elongation rate is an approximately linear function of $\hat{A}$:

$$v_{\text{el}} \approx k_r C_1(\hat{A} - 1).$$

4. When $n$ is high and $C_n \ll \hat{A}^n C_1$, the elongation rate is approximately a piecewise linear function:

$$v_{\text{el}} \approx \begin{cases} k_r C_n(\hat{A} - 1), & \hat{A} < 1, \\ k_r C_1(\hat{A} - 1), & \hat{A} \geq 1. \end{cases} \tag{S7}$$

Interestingly, this behaviour is analogous to rectified conductances seen in ion channels [2].

##### A.2.2 Bond graph component

To use the reduced model in a bond graph, the kinetic parameters in Eq. (S6) need to be expressed in terms of bond graph parameters. We note that since the kinetic parameters $k_f$ and $k_r$ are identical between elongation steps, $K_{C_{i+1}}/K_{C_i} = \alpha$ for some constant $\alpha$. We also note that from Eq. (1) in the main text, $K_i x_i = e^{\bar{\mu}_i}$. Then

$$k_r C_1 = r_n K_{C_n} x_{C_1} = r_n \frac{K_{C_n}}{K_{C_1}} K_{C_1} x_{C_1} = r_n \alpha^{n-1} e^{\bar{\mu}_{C_1}}, \tag{S8}$$

$$k_r C_n = r_n K_{C_n} x_{C_n} = r_n e^{\bar{\mu}_{C_n}}, \tag{S9}$$

$$\hat{A} = \frac{k_f A}{k_r} = \frac{r_i K_{C_i} K_A x_A}{r_i K_{C_{i+1}}} = e^{\bar{\mu}_A} \alpha^{-1}, \tag{S10}$$

where $r_i$ is the rate parameter for the reaction $C_{i-1} + A \rightleftharpoons C_i$.

Therefore, defining $m = n - 1$, we can express Eq. (S6) as

$$v_{\text{el}} = \left(r_n e^{\bar{\mu}_{C_1}} e^{m\bar{\mu}_A} - r_n e^{\bar{\mu}_{C_n}}\right) \frac{1 - e^{\bar{\mu}_A}\alpha^{-1}}{1 - e^{m\bar{\mu}_A}\alpha^{-m}} \tag{S11}$$

$$= r\left(e^{\bar{\mu}_{C_1}} - e^{\bar{\mu}_{C_n} - m\bar{\mu}_A}\right) \frac{1 - e^{\bar{\mu}_A}\alpha^{-1}}{(\alpha e^{-\bar{\mu}_A})^m - 1}, \tag{S12}$$

where $r = r_n \alpha^m$. Note that Eq. (S12) is a thermodynamically consistent expression as $v > 0$ only if $\mu_{C_1} + m\mu_A - \mu_{C_n} > 0$. Therefore, the reversible model of elongation can be encoded as a three-port bond graph component (Figure 4c in the main text) with the constitutive relations

$$\begin{aligned} v &= r\left(e^{\bar{\mu}_{C_1}} - e^{\bar{\mu}_{C_n} - m\bar{\mu}_A}\right) \frac{1 - e^{\bar{\mu}_A}\alpha^{-1}}{(\alpha e^{-\bar{\mu}_A})^m - 1}, \\ v_{C_1} &= -v, \\ v_{C_n} &= v, \\ v_A &= -mv. \end{aligned}$$

Thus, while the bond graph model of translation has been substantially reduced in complexity, it retains the thermodynamic consistency of the full model.

##### A.2.3 Lumped complex

The lumping scheme in § 2.4.2 of the main text groups several ribosomal complexes into the same species. Accordingly, the initial condition of the lumped complex $C_x$ is set to

$$C_x(t = 0) = \sum_{i=1}^{n-1} C_i(t = 0).$$

Assuming that $k_f A \gg k_r$, all complexes $C_i$ have similar concentrations at steady state, and hence

$$C_x \approx (n-1)C_i.$$

Due to the difference in amount between $C_x$ and $C_i$, the thermodynamic constant of $C_x$ is adjusted to maintain the same reaction rates between full and reduced models:

$$K_{C_x} = \frac{K_{C_1}}{n-1}.$$

##### A.2.4 Multiple lumped intermediates

As a generalisation to the earlier lumping scheme, one may lump complexes into more than one intermediate if greater accuracy is required. With $L$ lumps of length $k$, the reaction scheme simplifies to

$$M + R \underset{k_u}{\overset{k_b}{\rightleftharpoons}} C_0 \underset{k_r M}{\overset{k_f A}{\rightleftharpoons}} B_1 \overset{v_{\text{el},1}}{\rightleftharpoons} B_2 \overset{v_{\text{el},2}}{\rightleftharpoons} \ldots B_L \overset{v_{\text{el},l}}{\rightleftharpoons} C_n \underset{k_z}{\overset{k_t}{\rightleftharpoons}} R + P, \tag{S13}$$

where $B_i$ are groups of $k_i$ complexes and $v_{\text{el},i}$ is the net rate of the $i$-th lumped elongation reaction, following the rate law Eq. (S12) with $m = k_i$. The number of complexes in lump $B_i$ is

$$k_i = q + \mathbf{I}(i < r+1),$$

where $q = \lfloor (n-1)/L \rfloor$ is the integer quotient when $n-1$ is divided by $L$ with remainder $r = (n-1) - qL$, and $\mathbf{I}$ is the indicator function. This ensures consistency in the number of complexes (i.e. $\sum_{i=1}^{L} k_i = n-1$) while keeping the size of each lump as consistent as possible. A bond graph of this reduced model is shown in Figure 4d of the main text.

#### A.3 Degradation

In addition to transcription and translation, degradation processes are added for mRNA and proteins to avoid overaccumulation. In biology, both mRNA and proteins are known to be enzymatically degraded for regulatory purposes [3, 4]. Degradation rates can vary between mRNA and proteins, depending on their sequence, and are a major regulator of protein abundance and the timescale of their dynamics [5]. These involve the chemical transformation of such species into a sink $X$, representing the degradation products (e.g. nucleotides and amino acids). The degradation rate of a substrate $S$ is

$$v_{\text{deg},S} = r_{\text{deg},S}\left(e^{\bar{\mu}_S} - e^{\bar{\mu}_X}\right) = r_{\text{deg},S} K_S x_S - r_{\text{deg},S} e^{\bar{\mu}_X}.$$

While it is physically impossible for chemical potentials to be infinite, the chemical potential $\bar{\mu}_X$ is set to a very negative value so that

$$v_{\text{deg}} \approx r_{\text{deg},S} K_S x_S,$$

and hence the decay rate of the molecule is $r_{\text{deg},S} K_S$ [$\text{s}^{-1}$].

#### A.4 Gene expression model equations

The differential equations implied by the bond graph in Figure 5b of the main text are

$$\begin{aligned}
\frac{\mathrm{d}x_M}{\mathrm{d}t} &= v_{\mathrm{Tc}}(x_A, x_M, \mu_I) - v_b(x_M, x_R, x_{C_0}) - v_{\mathrm{deg},M}(x_M) + v_1(x_{C_0}, x_A, x_{C_x}, x_M),\\
\frac{\mathrm{d}x_R}{\mathrm{d}t} &= -v_b(x_M, x_R, x_{C_0}) + v_t(x_{C_n}, x_P, x_R),\\
\frac{\mathrm{d}x_P}{\mathrm{d}t} &= v_t(x_{C_n}, x_P, x_R) - v_{\mathrm{deg},P}(x_P),\\
\frac{\mathrm{d}x_{C_0}}{\mathrm{d}t} &= v_b(x_M, x_R, x_{C_0}) - v_1(x_{C_0}, x_A, x_{C_x}, x_M),\\
\frac{\mathrm{d}x_{C_x}}{\mathrm{d}t} &= v_1(x_{C_0}, x_A, x_{C_x}, x_M) - v_{\mathrm{el}}(x_{C_x}, x_{C_n}, x_A),\\
\frac{\mathrm{d}x_{C_n}}{\mathrm{d}t} &= v_{\mathrm{el}}(x_{C_x}, x_{C_n}, x_A) - v_t(x_{C_n}, x_P, x_R),
\end{aligned}$$

where the reaction fluxes are

$$v_{\mathrm{Tc}} = \bar{r}_{\mathrm{Tc}} \frac{K_A x_A - K_M x_M}{(1 + \frac{K_A x_A}{R_{bA}} + \frac{K_M x_M}{R_{bM}})(1 + \frac{e^{\bar{\mu}_I}}{R_{bI}})}, \tag{S14}$$

$$v_b = r_b(K_M K_R x_M x_R - K_{C_0} x_{C_0}), \tag{S15}$$

$$v_1 = r_1(K_{C_0} K_A x_{C_0} x_A - K_{C_x} K_M x_{C_x} x_M), \tag{S16}$$

$$v_{\mathrm{el}} = r_{\mathrm{el}} \left(K_{C_x} K_A^{(n-1)} x_{C_x} x_A^{(n-1)} - K_{C_n} x_{C_n}\right) \frac{1 - K_A x_A \alpha^{-1}}{1 - \left(\frac{K_A x_A}{\alpha}\right)^{n-1}}, \tag{S17}$$

$$v_t = r_t(K_{C_n} x_{C_n} - K_P K_R x_P x_R), \tag{S18}$$

$$v_{\mathrm{deg},M} = r_{\mathrm{deg},M} K_M x_M - r_{\mathrm{deg},M} e^{\bar{\mu}_{X,M}}, \tag{S19}$$

$$v_{\mathrm{deg},P} = r_{\mathrm{deg},P} K_P x_P - r_{\mathrm{deg},P} e^{\bar{\mu}_{X,P}}. \tag{S20}$$

#### A.5 Numerics

For some species, the thermodynamic parameter $K$ is larger than the maximum value permitted by 64-bit floats. However, these large parameters are counteracted by small reaction rates $r$ to result in moderate fluxes. To ensure that values remain numerically defined, some components use log-transformed parameters. In particular, the log-transformed species is defined by the equation

$$\bar{\mu} = \bar{\mu}^0 + \ln(x),$$

where $\bar{\mu}^0 = \ln(K)$; and log-transformed reactions are defined by the equation

$$v = e^{\bar{\mu}^a + \bar{\mu}^f} - e^{\bar{\mu}^a + \bar{\mu}^r},$$

where $\bar{\mu}^a = \ln(\kappa)$. Regulatory parameters such as $R_{bI}$ in Eq. (S2) can be log-transformed in a similar manner. This formulation is mathematically equivalent to the standard thermodynamic parameters, but avoids the need to evaluate large numbers.

### B Parameters

This section discusses how parameter values are chosen. Table S2 summarises the default values, with the justification discussed in later sections.

**Table S1: Default values used in gene expression model.**

| | Description | Default value | Section | References |
|---|---|---|---|---|
| $x_A$ | Number of energy molecules | $5.8 \times 10^6$ molecs | B.1 | [6, 7] |
| $x_M$ | Number of mRNA molecules | 10 molecs | B.1 | [8] |
| $x_R$ | Number of ribosomes | 5000 molecs | B.1 | [8, 9] |
| $n$ | ATP molecules required to synthesise protein | 1200 molecs | B.2 | [7] |
| $K_M$ | Thermodynamic parameter for mRNA | 1 $\text{molecs}^{-1}$ | B.2 | – |
| $K_R$ | Thermodynamic parameter for free ribosome | 1 $\text{molecs}^{-1}$ | B.2 | – |
| $K_A$ | Thermodynamic parameter for energy | 83.65 $\text{molecs}^{-1}$ | B.2 | [6, 10] |
| $K_{C_0}$ | Thermodynamic parameter for initial ribosomal complex | 1 $\text{molecs}^{-1}$ | B.2 | – |
| $\alpha$ | Ratio of parameters between ribosomal complexes | 3.49 | B.2 | [10] |
| $K_{C_1}$ | Thermodynamic parameter for ribosomal complex 1 | $\alpha$ $\text{molecs}^{-1}$ | B.2 | – |
| $K_{C_i}$ | Thermodynamic parameter for ribosomal complexes $(i > 1)$ | $\alpha^i$ $\text{molecs}^{-1}$ | B.2 | – |
| $K_P$ | Thermodynamic parameter for protein | $\alpha^n / e^{\bar{\mu}_{\text{termination}}}$ $\text{molecs}^{-1}$ | B.2 | [11] |
| $\bar{r}_{\text{Tc}}$ | Transcription rate | $1.13 \times 10^{-2}$ molecs/min | B.3.1 | [7] |
| $R_{bA}$ | Substrate binding parameter for transcription | 366.4 | B.3.1 | [7] |
| $R_{bM}$ | Product binding parameter for transcription | $10^6$ | B.3.1 | – |
| $R_{bI}$ | Inhibition parameter for transcription | $30K_P$ | B.3.1 | [7] |
| $r_b$ | Translation initiation rate | 0.001 molecs/min | B.3.2 | – |
| $r_1$ | Rate parameter for first elongation reaction | 1.04 molecs/min | B.3.2 | [12] |
| $r_i$ | Rate parameters for ith elongation reaction $(i > 1)$ | $5040/(e^{\bar{\mu}_A} K_{C_i})$ molecs/min | B.3.2 | [12] |
| $r_{\text{el}}$ | Rate parameter for elongation (reduced model) | $1.04 \times 10^{-5}$ molecs/min | B.3.2 | [12] |
| $r_t$ | Translation termination rate | $504000/K_{C_n}$ molecs/min | B.3.2 | – |
| $r_{\text{deg},M}$ | mRNA degradation rate | 0.1 molecs/min | B.3.3 | [7] |
| $r_{\text{deg},P}$ | Protein degradation rate | $0.02/K_P$ molecs/min | B.3.3 | [7] |
| $\bar{\mu}_{X,M}$ | Chemical potential of mRNA degradation products | $\ln(10^{-6})$ | B.3.3 | – |
| $\bar{\mu}_{X,P}$ | Chemical potential of protein degradation products | 0 | B.3.3 | – |

#### B.1 Molecule counts

**Energy**: Under nutrient-rich conditions, the ATP concentration has been estimated to be 9.63 mM in *E. coli* [6]. Assuming that the volume of an *E. coli* cell is 1 $\mu$m$^3$ [7], this corresponds to $5.8 \times 10^6$ molecules of ATP per cell. We use this as the reference number of energy molecules in a cell.

**mRNA**: In simulating translation in isolation, we assume there are 10 mRNA molecules per cell. This corresponds to a relatively high transcript count for a single gene [8].

**Ribosomes**: The number of ribosomes in an *E. coli* cell can vary between 6800–72000 [8]. However, the majority of ribosomes are occupied in translating proteins. At high growth rates, Dai et al. [9] estimated that 80% of ribosomes are actively translating protein. Thus, we assume there were 5000 free ribosomes per cell, corresponding to a total ribosome number of approximately 25000. Models are initialised with virtually all ribosomes in the free state; complex amounts are initialised to $10^{-6}$ molecules to avoid infinite potentials.

#### B.2 Thermodynamic parameters

**Reference species**: In the absence of free energy measurements, the mRNA ($M$) and ribosome ($R$) molecules are assumed to be reference species, so that their thermodynamic parameters are set to 1.

**Energy**: We assume that under the reference condition, the free energy of ATP hydrolysis is $-20k_bT$, corresponding to a Gibbs free energy of 49.5 kJ/mol at a temperature of 298 K. With a potential of $\bar{\mu}_A = 20$ and molecular count of $x_A = 5.8 \times 10^6$ molecules, the thermodynamic parameter is $K_A = 83.65$ molecules$^{-1}$ (Eq. (1) of the main text).

Our representation of ATP hydrolysis is a simplified version of the true biological reaction $\mathrm{ATP} + \mathrm{H_2O} \rightleftharpoons \mathrm{ADP} + \mathrm{Pi} + \mathrm{H^+}$. Thus, our model only considers ATP and neglects the other molecules. However, under conditions where the concentrations of the other molecules are constant, it can be shown that the equations reduce to our model.

**Complexes:** From Weiße et al. [7], the dissociation constant of the translation initiation reaction ($M + R \rightleftharpoons C_0$) is $K_b^d = K_{C_0}/(K_M K_R) = 1$. Then, since $K_M = K_R = 1$, it follows that $K_{C_0} = 1$.

We assume that the standard potentials of the chemical complexes differ by the energy required to form a peptide bond, or approximately $5k_BT$ of energy [10]. Since four ATP molecules are required to form a peptide bond, the standard potentials of successive complexes differ by $5/4k_BT$ and hence $\alpha = K_{C_{i+1}}/K_{C_i} = e^{5/4} = 3.49$. This corresponds to $\hat{A} = e^{\bar{\mu}_A - 5/4} = 1.38 \times 10^8$ under reference conditions (Eq. (S10)).

**Protein**: We assume that the termination of translation involves free energy contributions from protein folding and the dissociation of the protein synthesis complex. Because the free energy of protein folding ($\bar{\mu}_{\text{folding}} = -8.5$; Zeldovich et al. [11]) alone is insufficient to result in approximate irreversibility of termination, we assume a free energy of dissociation of $\bar{\mu}_{\text{dissociation}} = -11.5$ to result in a standard free energy of $\mu_{\text{termination}} = \bar{\mu}_{\text{folding}} + \bar{\mu}_{\text{dissociation}} = -20$. Hence, the thermodynamic parameter is $K_P = K_{C_n} e^{\bar{\mu}_{\text{termination}}} = \alpha^n e^{-20}$.

**Protein length:** We assume $n = 1200$, which corresponds to a protein length of 300 amino acids [7].

**Adjustment for mRNA copy number:** When simulating simulation in isolation, an adjustment is made to the above parameters. In considering the reverse reaction of $C_0 + A \underset{k_{r1}}{\overset{k_{f1}}{\rightleftharpoons}} C_1 + M$, a complex may only reoccupy the first site of its own transcript. To account for this, the reverse rate parameter $k_{r1}$ must be scaled down by the total number of mRNA molecules $M_{\text{tot}}$, i.e. $k_{r1} = k_r/M_{\text{tot}}$. Thus, $K_{C_1} = \alpha/(K_M M_{\text{tot}})$, $K_{C_i} = \alpha^i/(K_M M_{\text{tot}})$, $K_P = \alpha^n/(K_M M_{\text{tot}})$. However, since the number of mRNA molecules becomes variable once transcription and translation are coupled together, this adjustment is not applied under the coupled model.

#### B.3 Rate parameters

##### B.3.1 Transcription

Transcription parameters are calculated by matching enzyme kinetics behaviour to that of Weiße et al. [7]. The transcription rate law follows a Michaelis-Menten equation $v_{\text{Tc}} = wA/(A+\theta)$ with maximum flux $w = 4.14$ molecules/min and a half-saturation constant of $\theta = 4.38$ molecules. By comparing to Eq. (S14),

$$\begin{aligned} w/\theta &= \bar{r}_{\text{Tc}} K_A, \\ \theta &= R_{bA}/K_A. \end{aligned}$$

Solving the above equations, $\bar{r}_{\text{Tc}} = 1.13 \times 10^{-2}$ molecs/min and $R_{bA} = 366.4$. We assume $R_{bP} = 10^6$ so that the reverse reaction only saturates at high concentrations, and that $R_{bI} = 30e^{\bar{\mu}_I^0}$ to achieve inhibition at 30 molecules of inhibitor.

##### B.3.2 Translation

**Binding rate**: The binding rate constant is set to $r_b = 0.001$ to achieve biologically feasible protein copy numbers (1000's to 10000's) at steady-states with 10–100 mRNA molecules per cell [8].

**Elongation:** Under nutrient-rich conditions, the elongation rate has been measured to be 1260 amino acids per minute [12]. Since we model four elongation steps per amino acid, the forward elongation rate is $k_f A = 4 \times 1260 = 5040$ molecules/min. Then, since $k_f = r_i K_A K_{C_{i-1}}$ for $i = 1, \ldots, n$, $r_i = 5040/(K_A A)/K_{C_i} = 5040/(e^{20} K_{C_i})$. In the reduced model, these rate parameters correspond to $r_{\text{el}} = k_f A \alpha / e^{20}/K_{C_1} = 1.04 \times 10^{-5}$ molecules/min.

**Termination:** Termination is assumed to be fast relative to elongation, with a forward rate constant 100 times that of an elongation step. Thus, $r_t = 100 k_f A/K_{C_n} = 504000/K_{C_n}$.

##### B.3.3 Degradation

**mRNA degradation:** The mRNA degradation decay rate is set to 0.1 $\text{min}^{-1}$, in line with Weiße et al. [7]. The reaction rate associated with degradation is therefore set to $r_{\text{deg,M}} = 0.1/K_M = 0.1$ molecules/min. A negative degradation product potential of $\bar{\mu}_{X,M} = \ln(10^{-6})$ is used to approximate an irreversible reaction.

**Protein degradation:** The protein degradation decay rate is set to 0.02 $\text{min}^{-1}$, so that proteins degrade slower than mRNA. The reaction rate associated with degradation is therefore set to $r_{\text{deg,P}} = 0.02/K_P$ molecules/s. A degradation product potential of $\bar{\mu}_{X,P} = 0$ is used to approximate an irreversible reaction.

##### B.4 Toggle switch and repressilator

The following parameters are modified in simulating the toggle switch and repressilator:

- To increase expression levels, the binding constant is increased to $r_b = 0.01$ molecules/min
- The inhibition parameter is set to $R_I = (100K_P)^h$ so that inhibition occurs at protein concentrations on the order of 100 molecules Weiße et al. [7]
- The mRNA degradation rate is increased to $\log(2)/2$ min$^{-1}$, consistent with Weiße et al. [7]
- The protein degradation rate is increased to $\log(2)/4$ min$^{-1}$, consistent with Weiße et al. [7]

#### C Simplification under excess energy

##### C.1 Gene expression

Here we consider the reduced gene expression model with one lumped complex (§ 2.5 in the main text). Under conditions where $A$ is high, Eq. (S14) becomes

$$v_{\mathrm{Tc}} \approx v^*_{\mathrm{Tc}} = \frac{\bar{r}_{\mathrm{Tc}} R_{bA}}{1 + e^{\bar{\mu}_I}/R_{bI}}.$$

Also, due to the high amount of $A$, the elongation reactions can be assumed to be irreversible and instantaneous, so complexes are immediately moved to their terminal complex. Thus, the reaction network reduces to

$$\begin{aligned} \varnothing \to M &\to \varnothing \\ M + R &\to M + C_n \\ C_n &\rightleftharpoons R + P \\ P &\to \varnothing. \end{aligned}$$

For ease of analysis, we make the additional assumptions that (i) the mRNA transcription and degradation dynamics are faster than translation and (ii) termination is instantaneous and irreversible. Under these further assumptions, the reaction network becomes

$$\begin{aligned} \varnothing \to M &\to \varnothing \\ M + R &\to M + R + P \\ P &\to \varnothing, \end{aligned}$$

where the mRNA dynamics follows the equation

$$\begin{aligned} \frac{\mathrm{d}x_M}{\mathrm{d}t} &\approx v^*_{\mathrm{Tc}}(\bar{\mu}_I) - r_{\mathrm{deg},M} K_M x_M = 0 \\ \Rightarrow x_M &= \frac{v^*_{\mathrm{Tc}}(\bar{\mu}_I)}{r_{\mathrm{deg},M} K_M}. \end{aligned}$$

Using the steady-state value of $M$, the equation for protein dynamics becomes

$$\frac{\mathrm{d}x_P}{\mathrm{d}t} = r_b K_M K_R x_M x_R - r_{\mathrm{deg},P} K_P x_P = r_b K_M K_R x_R \frac{v^*_{\mathrm{Tc}}(\bar{\mu}_I)}{r_{\mathrm{deg},M} K_M} - r_{\mathrm{deg},P} K_P x_P.$$

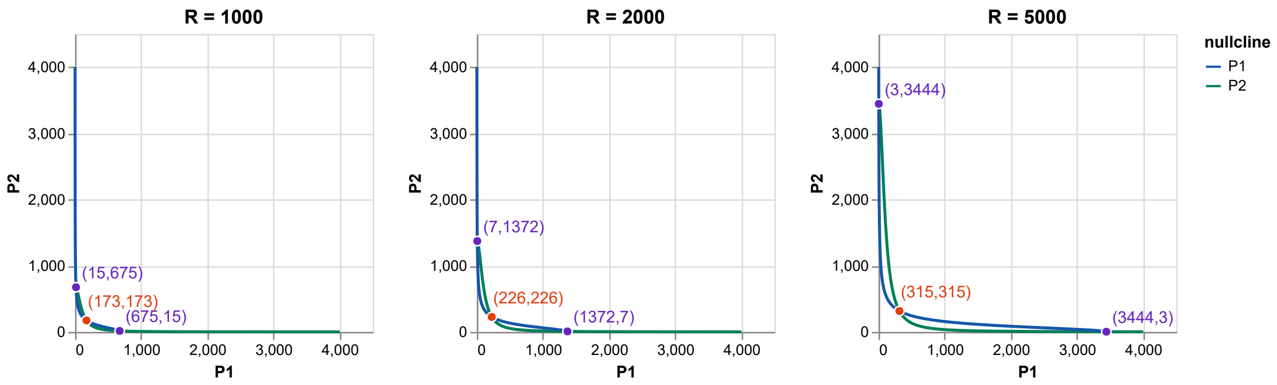

**Figure S1: A phase plane of the toggle switch.** Lines show the nullclines of the system. Purple points represent the stable fixed points, and the orange points are the unstable fixed points.

Thus, under the limit of high energy levels, the governing differential equations are of a similar form to previous works [13].

At steady state,

$$x_P = \frac{r_b K_R x_R v^*_{\mathrm{Tc}}(\bar{\mu}_I)}{r_{\mathrm{deg},M} r_{\mathrm{deg},P} K_P},$$

and the translation rate is the positive term of $\mathrm{d}x_P/\mathrm{d}t$:

$$v_{\mathrm{trans}} = \frac{r_b K_R x_R v^*_{\mathrm{Tc}}(\bar{\mu}_I)}{r_{\mathrm{deg},M}}.$$

#### C.2 Toggle switch

Under the above assumptions, the toggle switch follows the following equations at steady state:

$$\frac{\mathrm{d}x_{P1}}{\mathrm{d}t} = \frac{r_b K_M K_R x_R}{r_{\mathrm{deg},M} K_M} \frac{\bar{r}_{\mathrm{Tc}} R_{bA}}{1 + (K_P x_{P2})^2 / R_{bI}} - r_{\mathrm{deg},P} K_P x_{P1},$$
$$\frac{\mathrm{d}x_{P2}}{\mathrm{d}t} = \frac{r_b K_M K_R x_R}{r_{\mathrm{deg},M} K_M} \frac{\bar{r}_{\mathrm{Tc}} R_{bA}}{1 + (K_P x_{P1})^2 / R_{bI}} - r_{\mathrm{deg},P} K_P x_{P2}.$$

A phase plane of this system is shown in Figure S1, with the nullclines

$$x_{P1} = x_{P1}(x_{P2}) = \frac{r_b K_M K_R x_R}{r_{\mathrm{deg},M} K_M r_{\mathrm{deg},P} K_P} \frac{\bar{r}_{\mathrm{Tc}} R_{bA}}{1 + (K_P x_{P2})^2 / R_{bI}},$$
$$x_{P2} = x_{P2}(x_{P1}) = \frac{r_b K_M K_R x_R}{r_{\mathrm{deg},M} K_M r_{\mathrm{deg},P} K_P} \frac{\bar{r}_{\mathrm{Tc}} R_{bA}}{1 + (K_P x_{P1})^2 / R_{bI}}.$$

Thus, under the reference parameters, there are three fixed points, with two stable states. As the number of ribosomes decreases, the stable points move towards the origin, leading to a decrease in bistability index.

### D Energetic analysis

A strength of the bond graph approach is that it can reveal insights into the thermodynamic efficiency of biological processes. We examine the energetics of translation here. Translation

**Table S2: Energetic analysis of translation.** Models of translation with $n = 1200$ are simulated with different amounts of energy. The values recorded are the translation rate ($v_{\text{trans}}$ [molecules/s]), energy supplied ($\bar{\mu}_A$), chemical potential of protein ($\bar{\mu}_P$), thermodynamic efficiency ($\eta$), free energy of ribosomal binding ($\Delta G_{\text{binding}} = \bar{\mu}_{C_0} - \bar{\mu}_M - \bar{\mu}_R$), free energy of elongation ($\Delta G_{\text{elongation}} = \bar{\mu}_{C_n} + \bar{\mu}_M - \bar{\mu}_{C_0} - n\bar{\mu}_A$) and free energy of termination ($\Delta G_{\text{termination}} = \bar{\mu}_R + \bar{\mu}_P - \bar{\mu}_{C_n}$).

| $A$ | $v_{\text{trans}}$ | $\mu_A$ | $\mu_P$ | $\eta$ (%) | $\Delta G_{\text{binding}}$ | $\Delta G_{\text{elongation}}$ | $\Delta G_{\text{termination}}$ |
|---|---|---|---|---|---|---|---|
| $1 \times 10^5$ | 42.1 | 15.9 | 1486.9 | 7.77 | $-11.4$ | $-17629.2$ | $-9.16 \times 10^{-4}$ |
| $1 \times 10^6$ | 49.0 | 18.2 | 1486.9 | 6.79 | $-13.7$ | $-20390.0$ | $-9.57 \times 10^{-4}$ |
| $5.8 \times 10^6$ | 49.8 | 20.0 | 1486.9 | 6.20 | $-15.4$ | $-22497.7$ | $-9.61 \times 10^{-4}$ |

can be seen as the use of energy from ATP hydrolysis to form peptide bonds in proteins, with the overall reaction

$$nA \rightleftharpoons P.$$

Thus, we can define the thermodynamic efficiency $\eta$ of translation as the proportion of total energy from ATP hydrolysis used to form proteins:

$$\eta = \frac{\bar{\mu}_P}{\bar{\mu}_A}.$$

We simulate the translation model for different values of $A$ and calculate the thermodynamic efficiency for each of these energetic states (Table S2). The thermodynamic efficiency of translation is 6–8%, which is low compared to many other biological processes [14]. While this may be surprising at first glance, the apparently wasteful nature of translation has previously been attributed to kinetic proofreading, a mechanism for reducing the error rate [15]. While increasing $A$ increases the translation rate $v_t$, it also reduces the efficiency since the input potential $\mu_A$ increases without a corresponding increase in the output potential $\mu_P$.

The thermodynamic analysis in Table S2 also allows the thermodynamic consistency of the model to be verified. The free energies of binding ($M + R \rightleftharpoons C_0$), elongation ($C_0 + nA \rightleftharpoons C_n + M$) and termination ($C_n \rightleftharpoons R + P$) are defined as $\Delta G_{\text{binding}}$, $\Delta G_{\text{elongation}}$ and $\Delta G_{\text{termination}}$ respectively. One can easily verify that energy balance is satisfied, i.e. $\bar{\mu}_P - n\bar{\mu}_A = \Delta G_{\text{binding}} + \Delta G_{\text{elongation}} + \Delta G_{\text{termination}}$. The second law of thermodynamics is also satisfied since all the free energies are negative.

The free energies also provide a more detailed picture of how power is dissipated. The vast majority of energy is dissipated during elongation. While the binding free energy is much lower by comparison, it is still far from equilibrium. On the other hand, due to the fast nature of the termination reaction, it effectively operates at equilibrium ($\Delta G_{\text{elongation}} \approx 0$). Thus, it is likely that both the binding and elongation rates control the rate of protein translation, which is consistent with current hypotheses [16, 17].

### E Supplementary figures

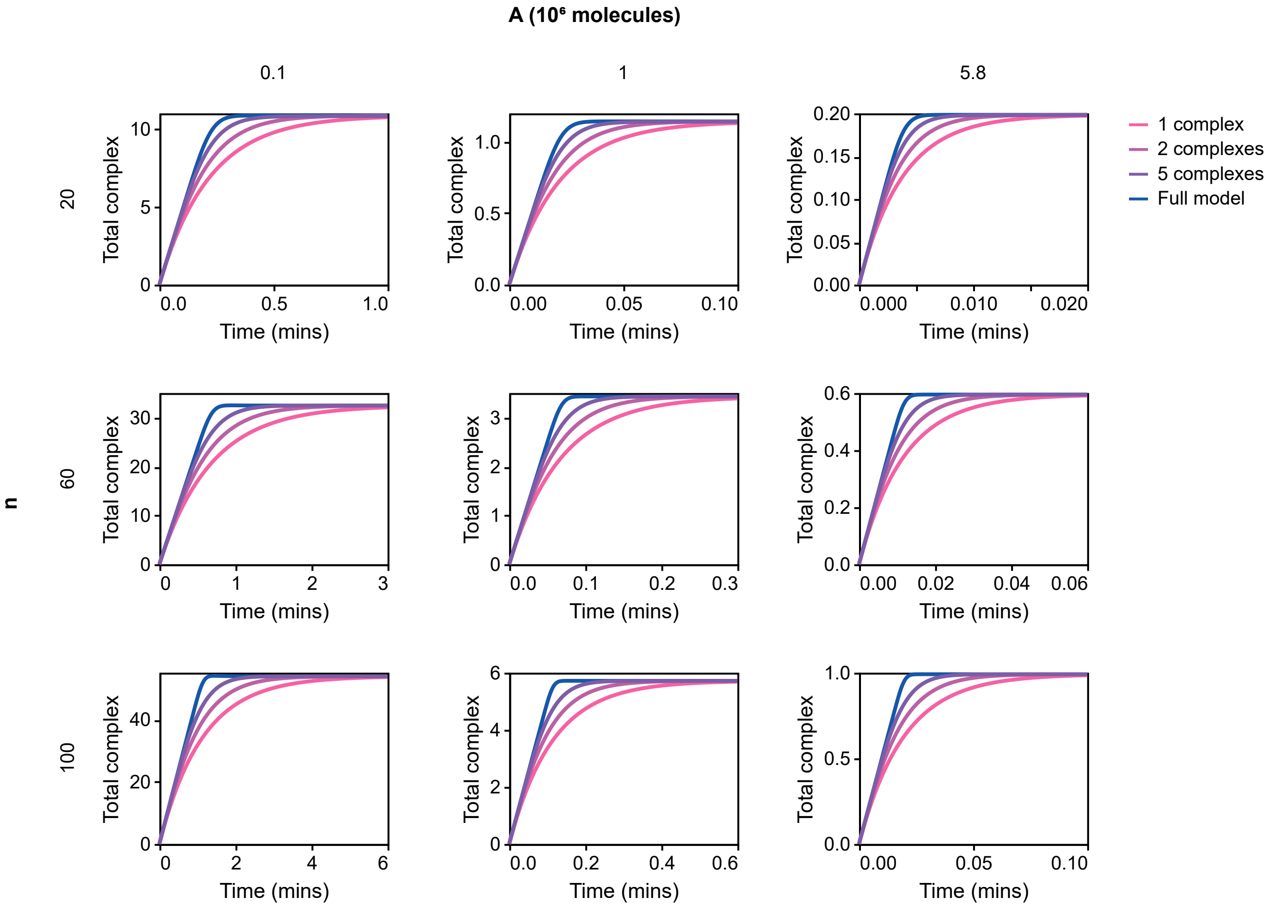

**Figure S2: Comparison of total complex amount for the full model.** The energetic state $A$ is varied horizontally, and the chain length $n$ is varied vertically. The translation rate is given in proteins per minute. Simulations are run with 10 mRNA molecules, 5000 ribosomes and $\bar{\mu_P} = -15$ to approximate the immediate removal of proteins.

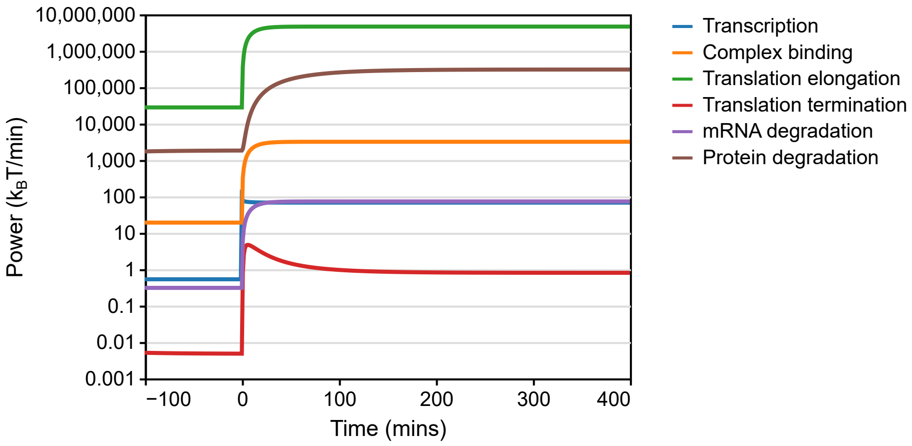

**Figure S3: Power consumption of gene regulation.** The gene expression model in Figure 10a–c of the main text is simulated, and the power dissipation rate of each process is plotted against time. The power dissipation rate of each reaction is defined as the sum of power through all ports of the corresponding reaction. The power disspation for elongation is defined as the sum of power dissipation rates for the first elongation reaction (Re:$r_1$) and the elongation module $\mathrm{Re}_{\mathrm{tl}}$.
